# Predicting long pendant edges in model phylogenies, with applications to biodiversity and tree inference

**DOI:** 10.1101/2021.09.11.459915

**Authors:** Sergey Bocharov, Simon Harris, Emma Kominek, Arne Ø. Mooers, Mike Steel

## Abstract

In the simplest phylogenetic diversification model (the pure-birth Yule process), lineages split independently at a constant rate λ for time *t*. The length of a randomly chosen edge (either interior or pendant) in the resulting tree has an expected value that rapidly converges to 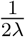 as *t* grows, and thus is essentially independent of *t*. However, the behaviour of the length *L* of the *longest* pendant edge reveals remarkably different behaviour: *L* converges to *t*/2 as the expected number of leaves grows. Extending this model to allow an extinction rate *μ* (where *μ* < λ), we also establish a similar result for birth-death trees, except that *t*/2 is replaced by *t*/2 · (1 – *μ*/λ). This ‘complete’ tree may contain subtrees that have died out before time *t*; for the ‘reduced tree’ that just involves the leaves present at time *t* and their direct ancestors, the longest pendant edge length *L* again converges to *t*/2. Thus, there is likely to be at least one extant species whose associated pendant branch attaches to the tree approximately half-way back in time to the origin of the entire clade. We also briefly consider the length of the shortest edges. Our results are relevant to phylogenetic diversity indices in biodiversity conservation, and to quantifying the length of aligned sequences required to correctly infer a tree. We compare our theoretical results with simulations, and with the branch lengths from a recent phylogenetic tree of all mammals.

## Introduction

Stochastic birth–death models (of speciation and extinction) model the tree-like diversification of species over evolutionary time-scales and play an important role in systematic biology. These models trace back to a seminal paper of Yule (1925), and a rich literature of probabilistic modelling of birth–death processes has developed, from the 1940s to the present. These stochastic models lead to predictions concerning the ‘shape’ of phylogenetic trees, and thereby allow the testing of different speciation-extinction models. The models also allow the formulation of estimators for speciation and extinction rates (based on a given model) from large phylogenies (see e.g. Nee et al. (1994)) and provide priors for Bayesian phylogenetic methods. Importantly, the birth–death process tends to produce trees whose distribution of splitting times is intermediate between the deep splits expected in adaptive radiations (Gavrilets and Vose, 2005) and the shallow splits expected under density-dependent dynamics (Hey, 1992). Although more recent compilations are required, published trees tend to produce mildly deeper-than-expected splitting times (Morlon et al., 2010).

Mathematical investigations into phylogenetic diversification models have also led to new insights and predictions (e.g. Aldous (2001); Aldous and Popovic (2005); Lambert and Stadler (2013), and, most recently, Louca and Pennell (2020)). This last paper established an inherent limitation on the extent to which it is possible to identify an unknown diversification model from an observed phylogeny, regardless of its size.

One aspect of any birth–death process is the length it predicts for edges of the tree (note that ‘edge’ is synonymous with ‘branch’ and a ‘pendant edge’ is one that is incident with a tip (or leaf) at the present). The simplest such model, the *Yule process*, has a single constant speciation rate λ and the expected length of a randomly chosen edge in a Yule tree grown for time *t* quickly converges to 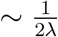 (Steel and Mooers, 2010) as *t* grows; the factor of 2 in the denominator may at first appear surprising but is due to the process taking place on a binary tree rather than along a path). Similar results for the length of a randomly selected edge are known for birth–death trees (Mooers et al., 2012; Stadler and Steel, 2012).

In this paper, we focus on the length of the *longest* pendant edge in a tree generated by a birth–death process, for both the complete tree (which is relevant to studies involving total sampling through time, as with certain viral data-sets) and for the ‘reduced tree’, which corresponds to phylogenies reconstructed from genomic data sampled from individual taxa at the present.

For Yule trees (i.e. a birth process where extinction is set to 0), Gascuel and Steel (2010) showed that the expected length of the longest edge does not converge to 0 as λ → ∞ (more precisely, the proof of their Proposition 2.2 showed that, with strictly positive probability, an edge of length at least *t*/4 exists). Here, we first establish a sharper result: the longest pendant edge for a large Yule tree grows linearly with *t* as the expected number of leaves grows, and it converges to *t*/2. In particular, it is independent of any fixed value of λ, in contrast to the ‘average’ pendant edge length, which essentially only depends on λ (and not *t*) for large trees.

We then extend this result to birth–death models (i.e. with nonzero extinction), for both the ‘complete tree’ and for the ‘reduced tree’. For complete birth–death trees, the term *t*/2 is replaced by a term that is smaller but still proportional to t, whereas for reduced trees, the term *t*/2 again surfaces as the appropriate limit. Thus, in trees inferred from data at the present, the models predict that there is likely to be at least one extant species whose associated pendant branch attaches to the tree approximately half-way back in time to the origin of the entire clade.

We compare our theoretical results with simulations of Yule, and birth-death trees (both complete and reduced) and then describe some implications of this result for systematic biology. First, we show that for large phylogenies generated under a birth-death model, the most ‘evolutionary distinct’ taxa are likely to be those at the end of the longest pendant edges. Second, we describe how our results bear on the question of how much genomic data (number of aligned sequence sites) are required in order to infer a fully resolved tree correctly.

Finally, we compare our expectations with recent inferred trees for 114 mammal families, and provide some concluding comments.

## The Yule process

We first consider a simple pure-birth model, referred to as the *Yule process*. This process begins with a single species at time 0, which persists as a single lineage for a random time *τ*, where *τ* is exponentially distributed with rate parameter λ (i.e. *τ* ~ Exp(λ)). At time *τ*, this lineage splits into two lineages, representing a speciation event. These lineages then evolve under the same process and independently of each other and of all the previous history (i.e. they each persist for an independent exponential time and are replaced by two lineages at that point, and so on). We denote the resulting Yule tree at time *t* by 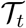, we let *L_t_* be the length of the longest pendant edge of 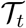 and let *S_t_* be the length of the shortest pendant edge of 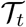, as indicated in Fig. 1.

**Fig. 1.**
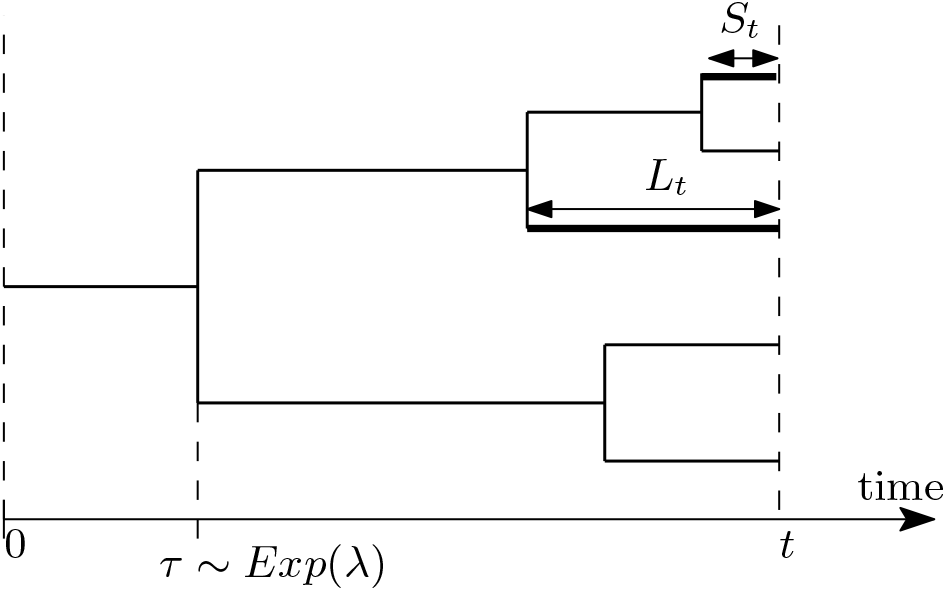
A Yule (pure-birth) tree starts with a single lineage at time *t* = 0, and each lineage persists for an exponentially distributed time *τ* with mean 1/λ. The four splitting events result in five leaf species at time *t*. The edge labels *L_t_* and *S_t_* denotes the length of the longest and shortest pendant edge of this tree, respectively.

### The longest pendant edge in a Yule (pure-birth) tree

#### Proposition 1

Let *L_t_* denote the length of the longest pendant edge in a Yule tree at time *t* with speciation rate λ. The following results hold:

i. 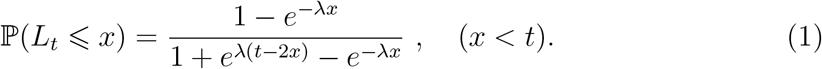
ii. The length of longest pendant edge when centred about *t*/2 (i.e., 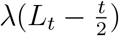) converges in distribution to a logistic distribution as λ*t* → ∞, where:

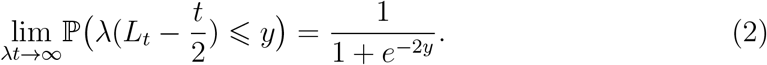
iii. Furthermore, *L_t_/t* converges to 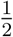 in mean and in probability as λ*t* becomes large (i.e. as λ*t* →∞, 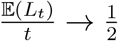, and for all *ϵ* > 0, 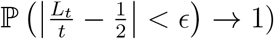.

Proposition 1 is a special case of the more general results stated later in Corollary 1 and Theorem 1 (the proofs for which are provided in the Appendix). For the Yule model, it is even possible to calculate the expected value of *L_t_* exactly, as we now show.

### The expected length of the longest pendant edge for Yule trees

For ease of presentation in what follows, let 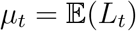, the expected length of the longest pendant edge in a Yule tree.

#### Proposition 2

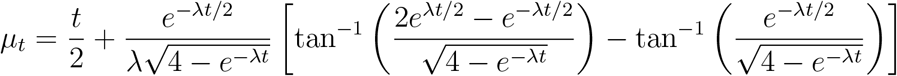

and 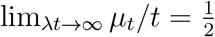.

#### Remarks

1. In Proposition 2, the (single) edge in a tree that has just one leaf is treated as a pendant edge. However, one might equally regard the stem edge as an interior edge, in which case a longest pendant edge would not exist in the one-leaf case (so one might then set *L_t_* = 0 in that case). Proposition 2 is easily adjusted to accommodate this. Let

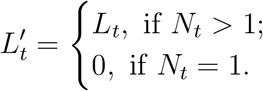 In contrast to *L_t_*, the probability distribution 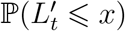 is now *continuous* at *x* = *t* (where it takes the value 1). It can be shown that 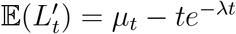 and that 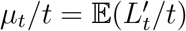 is monotone increasing, from 0 (at the limit *t* → 0^+^) to 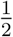 (as λ*t* → ∞). By contrast, 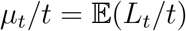 is monotone decreasing, from 1 (at the limit *t* → 0^+^) to 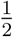 (as λ*t* → ∞).
2. *The shortest pendant branch length*. The distribution of the length of the shortest pendant edge (we denote this by *S_t_*) can also be exactly described. The condition that the shortest edge is longer than *x* is equivalent to the condition that each lineage present at time *t* – *x* does not split over the last time *x* (these independent events each have probability *e*^−λ*x*^). Since *N*_*t–x*_ ~ Geom(*e*^−λ(*t–x*)^), we find, for *x* ∈ [0, *t*]:

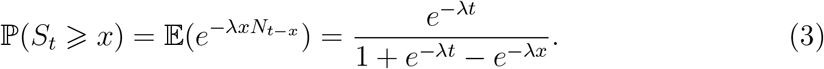 In particular, since *S_t_* ∈ [0, *t*], we have 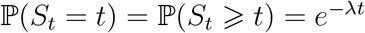, corresponding to no death of the initial ancestor (and 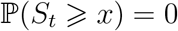 for *x* > *t*) thus:

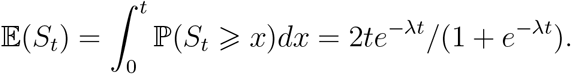 Thus, as λ*t* grows, 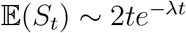. On the other hand, putting *x* = *ye*^−λ*t*^/λ in Eqn. (3) and noting that 1 – *e^−w^* ~ *w* for *w* small, we see that λ*e*^λ*t*^*S_t_* converges in distribution, with 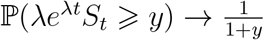 as λ*t* → ∞. This further implies that the ratio 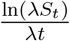 converges in probability to –1 as λ*t* grows.

## Long pendant edges in a (complete) birth–death phylogeny

The simple birth–death process generalises the Yule model from the previous section by allowing the extinction of lineages. In this model, each lineage (i.e. branch) present at any given time behaves independently, being replaced by two new lineages at a constant rate λ, and dying at a constant rate *μ*. Thus, each lineage persists for an independent random time *τ* which is exponentially distributed with rate λ + *μ*. At the end of its lifetime, it is either replaced by two new lineages, with probability λ/(λ + *μ*), or by none, with probability *μ*/(λ + *μ*). Lineages once alive behave independently of each other and of all the previous history, and in the same probabilistic manner as the parent (i.e. after an independent exponential time of rate λ + *μ*, it will terminate and either be replaced by 2 or 0 lineages, etc.)

In this section, we will extend our results for the Yule process to obtain results for the longest pendant edges in a birth–death tree. In the Section *‘Length of pendant edges in the reduced tree’*, we will then consider the pendant edges in the *reduced tree*, which is formed by tracing back the lineages that are alive at some fixed time *t* (i.e. we prune any lineages (subtrees) that have already died out before time *t*).

We start by recalling some well-known properties about birth–death processes.

### Some properties of birth–death trees

The following results concerning birth-death processes are classical (see e.g. Kendall (1948) or Grimmett and Stirzaker (2001)). Recall that the probability-generating function of a discrete random variable *X* is the function 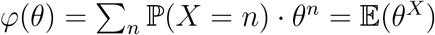 where *θ* is a formal variable and *n* ranges over all possible values *X* can take.

#### Lemma 1

The probability-generating function *φ_t_*(*θ*) of *N_t_* (the number of leaves alive at time *t* in the complete tree 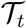) when λ > *μ* is given by:

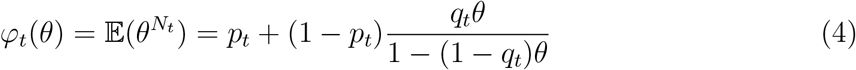

where:

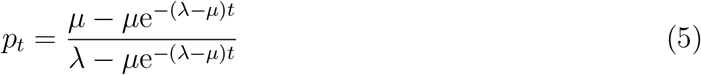

and

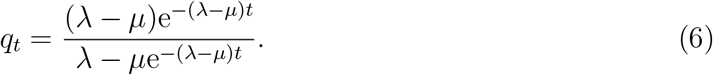

In particular, the probability that the process becomes extinct before time *t* is:

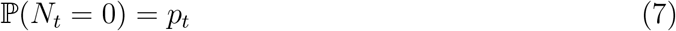

and, conditional on non-extinction by time *t*, *N_t_* has a geometric distribution on {1, 2, 3, · · · } with parameter *q_t_*. Formally:

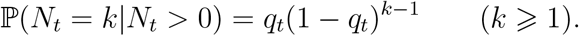

*N_t_* is said to have a *modified geometric* distribution written as *N_t_* ~ ModGeom(*p_t_, q_t_*).

Throughout this paper, we will let 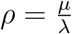, which is sometimes called the ‘turnover rate’ in phylogenetic diversification models and plays a fundamental role. For example, in the *supercritical* case (λ > *μ*), it follows from the formulae above (Eqns. (5) and (7)), that

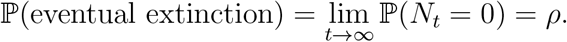

We assume throughout this paper that λ > *μ* ⩾ 0, and so 0 ⩽ *ρ* < 1. Thus the birth–death process is supercritical with a strictly positive probability of 1 – *ρ* of surviving forever.

### Length of pendant edges in complete trees

For any *t* ⩾ *x* ⩾ 0, define the random variable 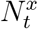 as follows:

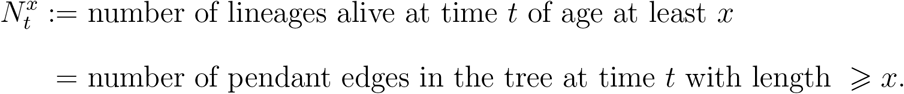

For the birth-death tree, we can find the distribution for the number of pendant edges greater than length *x* at time *t* explicitly. We will then be easily able to deduce the distributions for the sizes of the longest pendant edges at time *t*. For *x* ∈ [0, *t*], let:

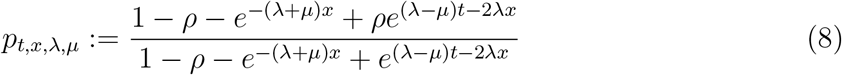

and

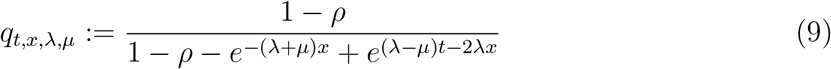

The proof of the following fundamental result is provided in the Appendix.

#### Lemma 2 (Distribution of the number of long pendant edges in a birth–death process)

The number of pendant edges of length at least *x* at time *t*, 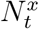, has a modified geometric distribution with 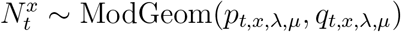. Formally:

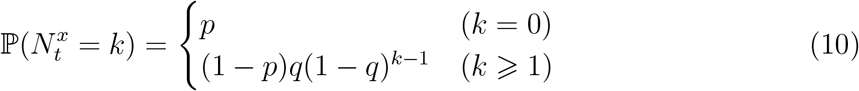

where *p* = *p*_*t,x*,λ, *μ*_ and *q* = *q*_*t,x*,λ, *μ*_, as given in Eqns. (8) and (9). In particular,

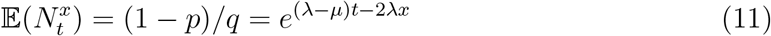

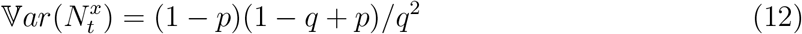

Let *L_t_* denote the length of the longest pendant edge that is still alive at time *t* in a birth–death tree at time *t*.

#### Corollary 1 (Distribution of the longest pendant edges in a birth–death process)

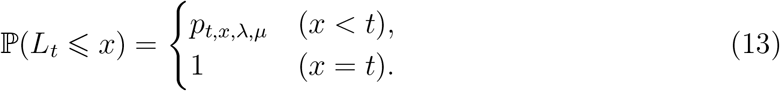

In particular, 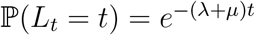, which corresponds to the initial ancestor neither branching nor dying over the entire time period *t*. Moreover, if 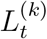 denotes the length of the *k*^th^ longest pendant edge at time *t*, then:

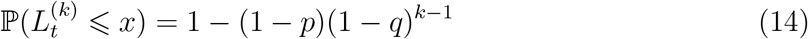

where *p* = *p*_*t,x*,λ, *μ*_ and *q* = *q*_*t,x*,λ, *μ*_, as given in Eqns. (8) and (9).

*Proof of Corollary 1*. This follows directly from the distribution of 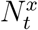 given in Lemma 2 since 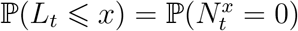 and 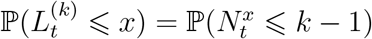.

**Remark:** There may be extinction by time *t* in the birth-death process when *μ* > 0, in which case *L_t_* = 0 and this event has probability:

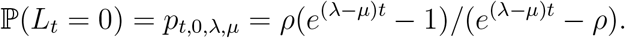

Similarly, there may not always be a *k*^th^ longest pendant edge (even for the Yule process), as only a finite number of lineages are present at time *t*. In such cases, by convention, we set the random variable 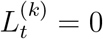 whenever *N_t_* < *k*, and we note that 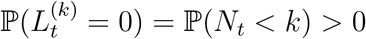 whenever *k* ⩾ 2.

### Distribution of the longest pendant edge in large complete trees

We now consider what happens for ‘large’ birth-death trees; by ‘large’, we mean the expected number of leaves in a birth-death tree (i.e. *e*^(λ–*μ*)*t*^) is large. If we regard *ρ* (= *μ*/λ) as a fixed value, then the expected number of leaves in a birth tree grows as a function of λ*t*. Thus, in the following theorem, we consider what happens as λ*t* grows (note that this may be due to λ becoming large even if *t* is not (for instance, a rapid species radiation in short time) or due to *t* becoming large (for instance, a tree that traces back deep into the past).

In this subsection, we assume that the birth-death process is supercritical with λ > *μ* and *ρ* is fixed. The expected number of leaves in the tree at time *t* can then be written as *e*^(1–*ρ*)λ*t*^, which grows as a function of λ*t*. Indeed, the number of leaves conditional on survival will also grow asymptotically at the rate *e*^(1–*ρ*)λ*t*^ whenever λ*t* → ∞. We can now state our main result for large complete birth-death trees, the formal proof of which is provided in the Appendix.

#### Theorem 1 (Longest pendant edges in the birth–death process as λ*t* grows)

Let λ > *μ* with *ρ* ≔ *μ*/λ being fixed. Conditional on the survival of the tree at time *t*, the following results hold:

i. For any 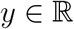, the number of pendant edges at time *t* that are longer than *t*(1 – *ρ*)/2 + *y*/λ converges in distribution as λ*t* → ∞ to a geometric distribution supported on {0,1, 2, … } as given by:

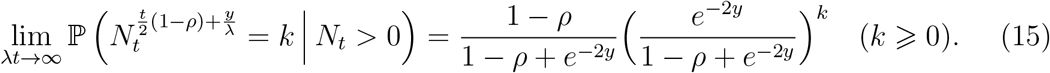
ii. In particular, the length of longest pendant edge when centred about *t*(1 – *ρ*)/2 (i.e. 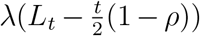) converges in distribution as λ*t* → ∞ to a logistic distribution, where

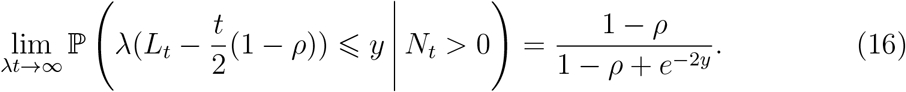 Furthermore, given survival up until time *t*, *L_t_*/*t* converges in probability to the constant (1 – *ρ*)/2, and in addition the average value of *L_t_*/*t* also tends to (1 – *ρ*)/2 (ie. 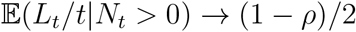).
iii. More generally, for each *k* ⩾ 1, the length of the *k*^th^ longest pendant edge when centred about *t*(1 – *ρ*)/2, that is, 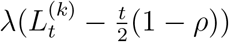, converges in distribution for large trees where

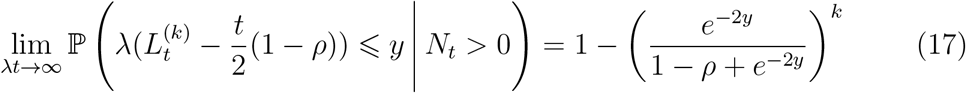 Furthermore, given survival up until time *t*, 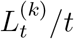 converges in probability to the constant (1 – *ρ*)/2, and in addition the average value of 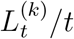 also tends to (1 – *ρ*)/2 (ie. 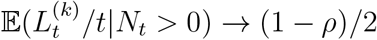).

## Length of pendant edges in the reduced tree

We now turn from complete trees to reduced trees based just on the leaves extant at the present time *t*. For this latter class of trees, we will show that the length of the longest pendant edge turns out to be (asymptotically) independent of the rates λ and *μ*, in contrast to the situation with complete trees, for which the asymptotic value was shown to be *t*/2 · (1 – *ρ*), and also in contrast to a randomly selected pendant edge from a reduced tree, for which the expected length does depend on *ρ* (Stadler and Steel, 2012).

More precisely, the *reduced tree* of the birth-death process, is the genealogical tree constructed only from the ancestors of the lineages still alive at time *t*. In other words, starting with the complete birth-death tree, we prune away any subtrees that died out before time *t* (or, equivalently, only keeping a lineage alive at time *s* < *t* if it has at least one descendant still alive at time *t*). This is illustrated in Fig. 2. The reduced tree is essentially the same as what is referred to as the ‘reconstructed tree’, except that in the latter the stem edge is often not present.

**Fig. 2.**
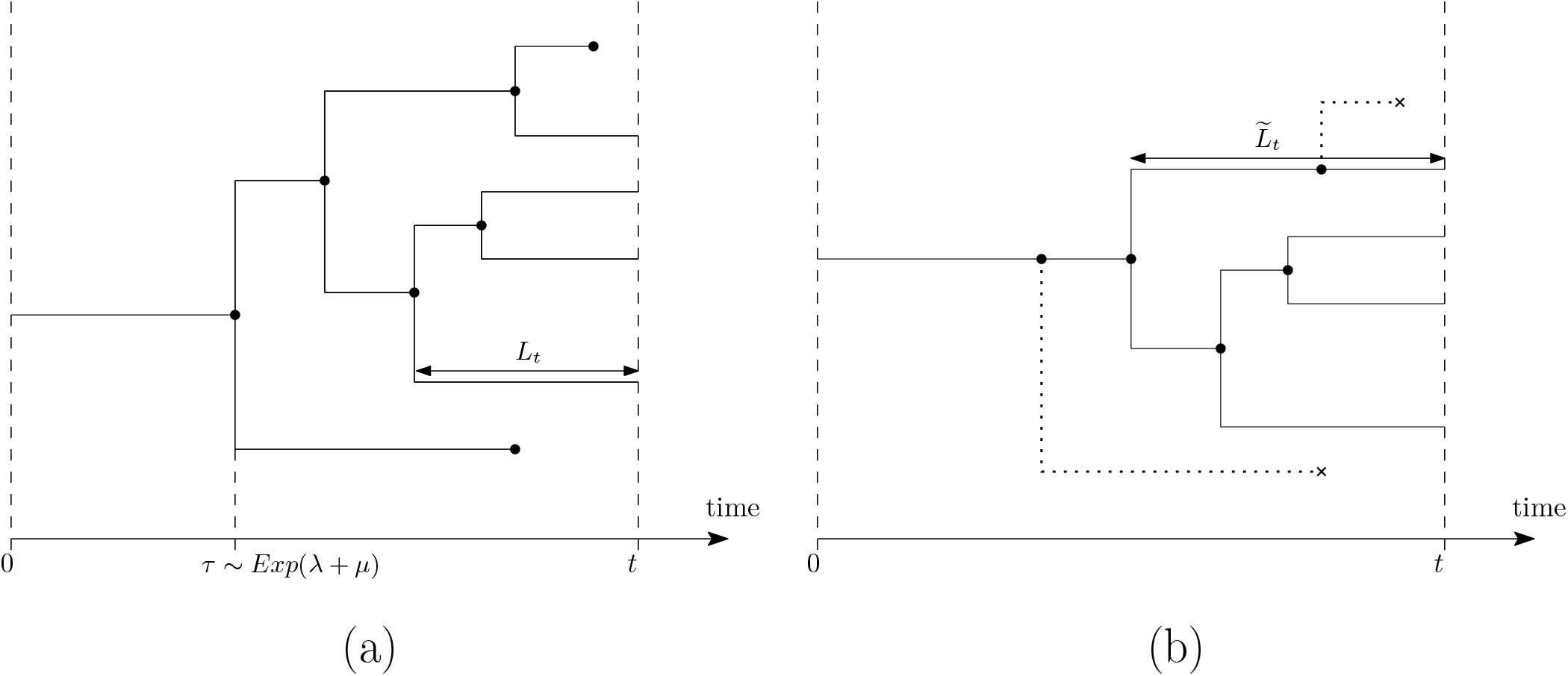
(a) A complete birth–death tree and (b) the corresponding reduced tree, where the dashed lineages have been pruned off. In (a), the edge of length *L_t_* is the longest pendant edge that is still alive at time *t* (the longer pendant edge below it dies before time *t*). Notice also that the length of the longest pendant edge in the reduced tree 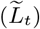 is greater than the length of the longest pendant edge of the complete tree (*L_t_*).

As any branches ending in a death are pruned away, a reduced tree for the birth-death process looks similar to a Yule tree. However, the probabilistic behaviour of the reduced tree is more complicated than that of a Yule tree. In a Yule tree, each individual branches into two at some constant rate λ. In the reduced tree, however, the rate of branching of each individual becomes time-dependant at rate λ(1 – *p_t–s_*) for any lineage alive at time *s*, where 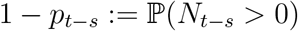 corresponds to the probability that at least one lineage born at time *s* will still be alive at time *t*.

Although the number of pendant edges that persist to time *t* remains unchanged as we prune away any sub-trees that died out in the birth-death tree, it is straightforward to see that these pendant edges can either remain the same or increase their length in the reduced tree (see Fig. 2). As a consequence, the behaviour of the long pendant edges in the reduced tree turns out to be somewhat different than in the corresponding complete birth-death tree when extinctions can occur (*μ* > 0).

### Length of pendant edges in a reduced tree

#### Lemma 3 (Distribution for the number of long pendant edges in the reduced birth-death tree)

The number of pendant edges of length at least *x* at time *t*, 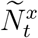, has a modified geometric distribution with 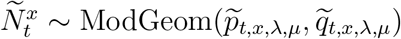. Formally:

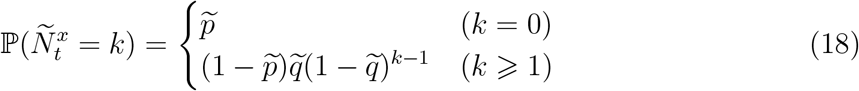

where 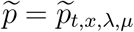 and 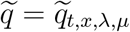 are given in the Appendix as Eqns. ((37)) and ((38)). In particular,

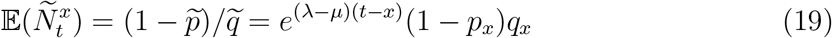

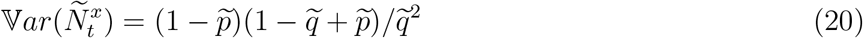

Let 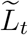 denote the length of the longest pendant edge in a reduced birth-death tree at time *t*.

#### Corollary 2 (Distribution of the longest pendant edges in the reduced birth–death tree for fixed values of λ and *t*)

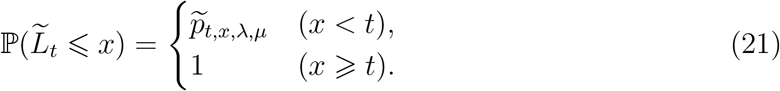

In particular, 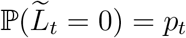, which corresponds to extinction by time *t*, and 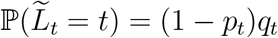, which corresponds to the initial ancestor having exactly one descendant alive at time *t*.

Moreover, if 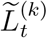 is the length of the *k*^th^ longest pendant edge at time *t*, then:

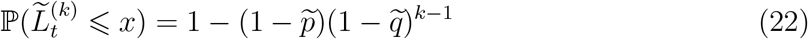

where 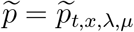 and 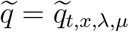, as in Eqns. (37) and (38).

### Length of the longest pendant edges in large reduced trees

When λ > *μ* > 0, conditional on survival, the reduced tree will become an increasingly large tree as λ*t* grows. Note that from Eqns. (5) and (6), as λ*t* → ∞:

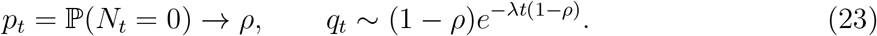

Using the formulae for 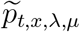 and 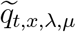 given by Eqns. (37) and (38) respectively in the Appendix, and setting 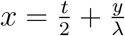, in those equations, we find that, as λ*t* → ∞,

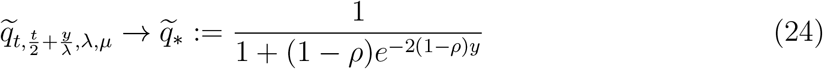

and

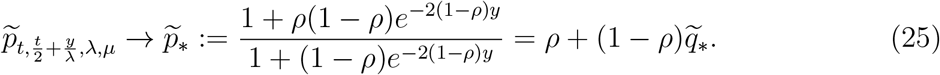

#### Theorem 2 (Longest pendant edges in the reduced tree as λ*t* grows)

Let λ > *μ* with *ρ* ≔ *μ*/λ being fixed. Conditional on the survival of the tree at time *t*, the following results hold:

i. For any 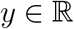, the number of pendant edges in the reduced tree at time *t* that are longer than *t*/2 + *y*/λ converges in distribution as λ*t* → ∞ to a geometric distribution supported on {0,1, 2,… } as given by:

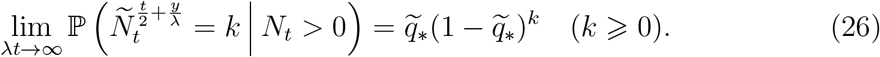
ii. In particular, the length of the longest pendant edge in the reduced tree when centred about *t*/2 (i.e. 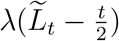) converges in distribution as λ*t* → ∞ to a logistic distribution, where

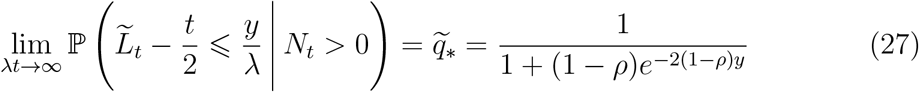 Furthermore, given survival up until time *t*, 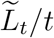 converges in probability to the constant 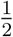, and in addition the average value of 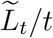 also tends to 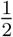 (ie. 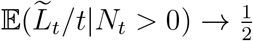.
iii. More generally, for each *k* ⩾ 1, the length of the *k*^th^ longest pendant edge in the reduced tree when centred about *t*/2 (i.e. 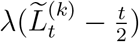) converges in distribution for large trees, where:

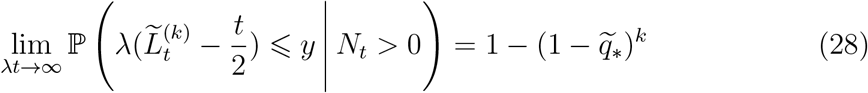 Furthermore, given survival up until time *t*, 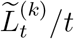 converges in probability to the constant 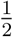, and in addition the average value of 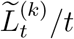 also tends to 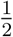 (ie. 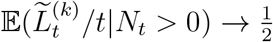.

Initially, it may seem surprising that the longest pendant edge in a large complete birth–death tree is roughly *t*(1 – *ρ*)/2 and this changes to roughly *t*/2 for the reduced tree, just like for Yule trees. This is because, if we condition a birth–death process to survive, its reduced tree will look very close to being a Yule tree with a branching rate λ(1 – *ρ*), at least until near to the end time *t* (in the reduced tree, lineages undergo binary branching at a rate of λ(1 – *p*_t–s_) at time *s*, but *p_t–s_* → *ρ* whenever λ(*t* – *s*) → ∞).

### Sampling at the present

Suppose that, in addition to a birth-death process (with rates λ and *μ* respectively), a proportion *σ* of the leaves are randomly sampled at the present. From Stadler (2009), the reduced tree on this pruned leaf set has the same distribution as a reduced birth-death tree with modified birth and death rates, as given by the following relationships:

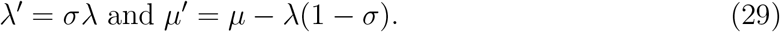

In order that *μ*′ ⩾ 0 one requires that *μ* ⩾ λ(1 – *σ*). Thus the results on long edges in birth-death trees can be extended to certain settings that involve sampling at the present.

## Simulation results

We used the *Treesim R* package (Stadler (2011); sim.bd.age function) to simulate 500 Yule trees (with λ = 0.416, *μ* = 0, *t* = 10) giving a target size of *n* = 64 extant species. We also simulated 500 complete birth-death trees (with λ = 1.11, *μ* = 0.5λ = 0.555, *t* = 10) retaining only trees with 1 or more extant species, giving a target size of *n* = 513 (since Nt, conditioned on non-extinction, has a geometric distribution with expected value 1/*q_t_*). We created a third set of 500 reduced birth-death trees by pruning all extinct species from the second set of complete birth-death trees using the *geiger* package (Pennell et al. (2014); drop.tip function).

We then calculated pendant edge lengths for all trees in each set using the ape, picante, phytools and geiger packages (Kembel et al., 2010; Revell, 2012; Pennell et al., 2014; Paradis and Schliep, 2019) and identified the longest pendant edge for each tree (denoted in this section by 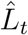). These observed 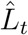 values were then compared to the theoretical expected values of *L_t_/t* (namely, 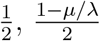 and 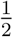) for the Yule, complete and reduced birth-death trees, respectively. The results are presented in Fig. 3.

**Fig. 3.**
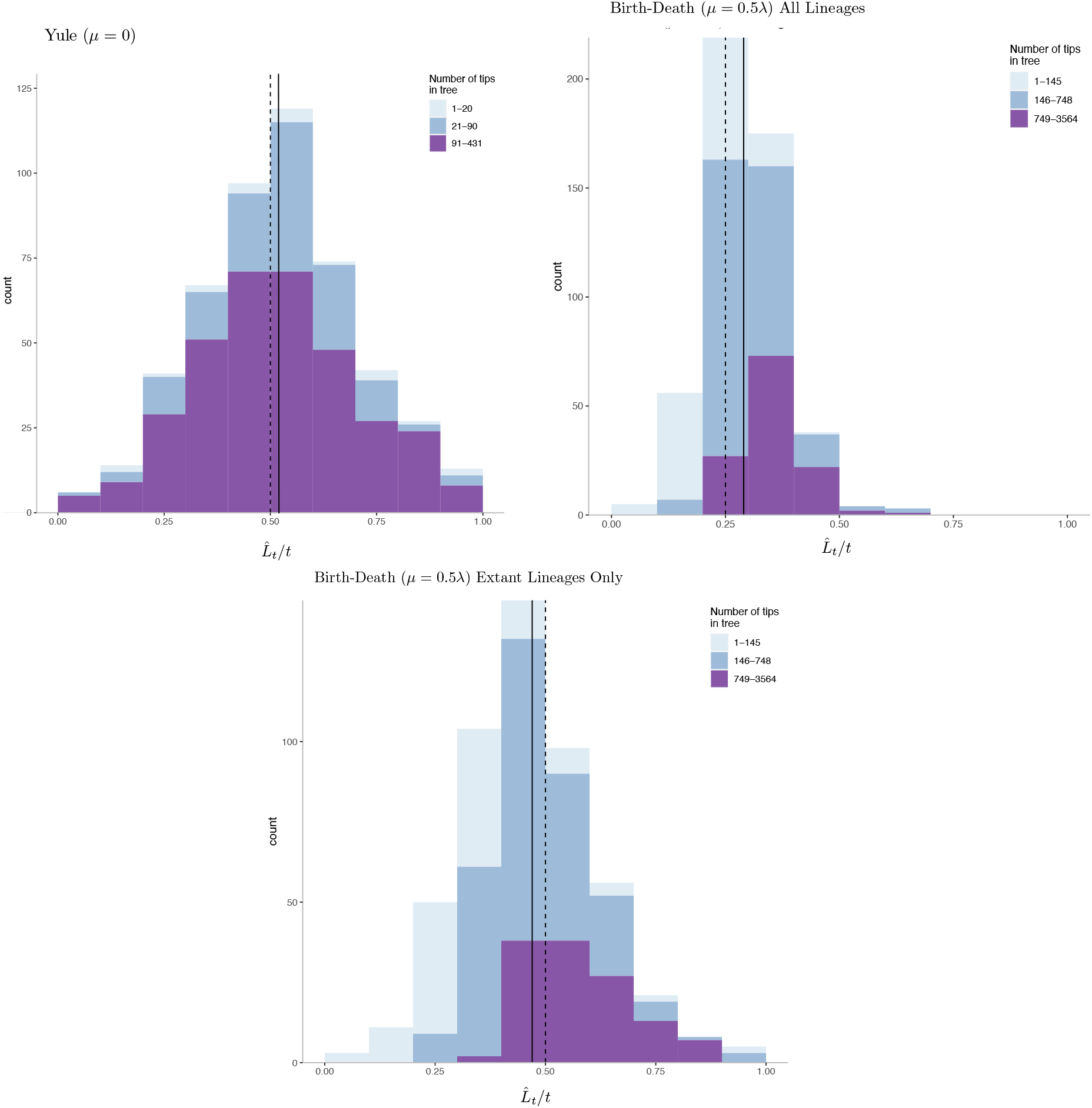
The longest pendant edges (that are still alive at time *t*) on simulated birth-death trees, with 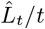 on the horizontal axis, and with trees sorted into three bins of small, medium and large trees. *Top Left*: The longest pendant edges (as 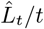) for 500 Yule trees (λ = 0.416, *μ* = 0) and depth *t* = 10. The dotted vertical line at 0.5 indicates the expected value of 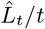 in the large tree limit (i.e. as λ*t* → ∞). Top *Right*: 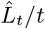 for complete birth-death trees (λ = 1.11, *μ* = 0.5λ = 0.555, *t* = 10). The dotted vertical line at 0.25 (= (1 – *ρ*)/2) indicates the expected value of 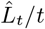 in the large tree limit and the solid line at 0.29 is the average value of 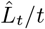. *Bottom*: 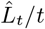 for reduced birth-death trees (λ = 1.11, *μ* = 0.555, *t* = 10). The dotted vertical line at 0.5 indicates the expected value of 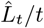 in the large tree limit and the solid line at 0.47 indicates the average value of 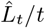.

Overall, 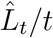 values are large, and in line with the theoretical predictions (note that the dashed lines refer to expected values of 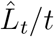 in the limit as the expected size of the trees tends to infinity (which, for *ρ* fixed, is equivalent to the limit as λ*t* → ∞)). Moreover, larger trees (having more leaves than expected) tended to have even longer-than-expected longest pendant edges, especially for the complete trees with extinction. This observation may reflect the “push of the past” phenomenon (Phillimore and Price, 2008) whereby many early splits occur in clades that persist and become large, a phenomenon that will both produce short stem edges and more room for an early lineage to persist to become a pendant edge. To the extent that the push of the past is real, this might make our expectations somewhat conservative (though see below for data from mammal families).

## Relevance to phylogenetic diversity

In biodiversity conservation, given a phylogenetic tree on a leaf set *X* of extant species, the *Fair Proportion (FP) index* (often called the ‘evolutionary distinctiveness’ index) is a way to assign the total sum of edge lengths across the tree ‘fairly’ to each of the extant species (Isaac et al., 2007; Redding, 2003; Redding et al., 2008). More precisely, suppose we have a phylogenetic tree *T* on the leaf set *X*, with edge lengths. We will assume (as in the rest of this paper and in most applications of FP), that these edge lengths correspond to (or are proportional to) time, and thus the sum of the lengths from the root to each leaf is the same (the so-called ‘ultrametric’ condition).

For a leaf *x* of *T*, let 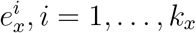 denote the edges on the directed path from *x* back to the root, let 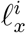 be the length of edge 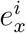, and let *n_i_*(*x*) be the number of leaves of *T* that are separated from the root of *T* by 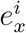 (i.e. the number of leaves of *T* descended from 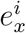). For a leaf *x* of a phylogenetic tree *T* with edge lengths ℓ, the FP index of *x*, denoted *FP*_(*T*,ℓ)_(*x*) (or, more briefly, *FP*(*x*)), is then defined as follows:

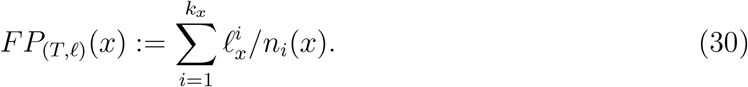

Since *n*_1_(*x*) is always equal to 1, we can re-write Eqn. (30) as follows:

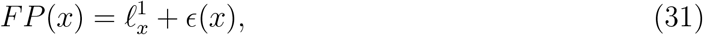

where:

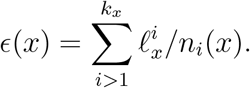

Two key features of the FP index are that (i) summing *FP*(*x*) over all the species *x* in *X* gives the total sum of edge lengths of the tree, and (ii) *FP*(*x*) has an equivalent description in terms of the Shapley value in cooperative game theory (Fuchs and Jin, 2015).

Fair proportion is a measure of evolutionary non-redundancy. More precisely, it provides a measure of the extent of sharing the products of evolution across species (a more formal justification of this statement under a model in which features arise at most once and are retained is described in Wicke et al. (2021)). An early empirical observation from (Redding et al., 2008) is that, on average, 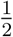 of the value of FP comes from the pendant edge (i.e.the value 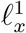) for Yule trees. Thus, species on very long pendant edges would be expected to be those with more truly unique features.

We can formalise this earlier empirical finding as follows. For a Yule tree *T* grown for time *t* at rate λ and a leaf *x* selected uniformly at random from those present at time *t* the following equation holds:

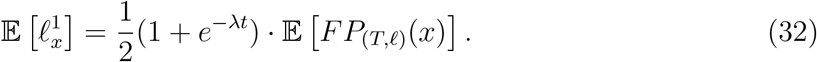

In particular, as λ*t* grows, the ratio of the expected pendant edge length to expected FP value for leaf *x* (i.e. 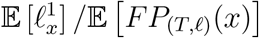) converges rapidly towards 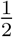. The proof of Eqn. (32) is provided in the Appendix.

We now present a result that states, roughly speaking, that in large reduced birth–death trees with λ > *μ*, a species with the highest FP score is expected to lie at the end of a pendant edge that has length close to *t*/2. One can easily construct trees where this expectation does not hold, so this prediction is not for trees in general, but rather for trees that have shapes captured by the birth–death model. The proof of the following theorem is provided in the Appendix.

### Theorem 3

For any *ϵ* > 0, and λ, *μ* fixed (λ > *μ*), consider a reduced birth-death tree *T* grown for time *t*. Then with probability tending to 1 as λ*t* grows (with *ρ* = *μ*/λ fixed), any species *x* that maximises *FP*(*x*) is the endpoint of a pendant edge of length at least 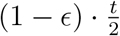.

### Mammal families conform to edge length predictions

We identified the longest pendant edges across 114 mammal families with ⩾ 3 species (Burgin et al., 2018; Upham et al., 2018). A cursory look suggests that several of the results above are predictive for empirical trees.

Firstly, the length of the longest pendant edge is a significant proportion of *t* (the time back to the origin of the clade) across a wide range of tree sizes, as summarised in Table 1 and the accompanying histogram. The values of 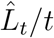 are less than 0.5 but appear to be increasing toward this value with increasing tree sizes, and the variation in 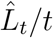 is also declining with tree sizes, both of which are consistent with Theorem 2.

**Table 1.** The ratio 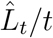 across phylogenetic trees for 114 mammal families

| Number of tips | Number of trees | mean of $\hat{L}_t/t$ | median of $\hat{L}_t/t$ | coefficient of variation of $\hat{L}_t/t$ |
| --- | --- | --- | --- | --- |
| 3 – 7 species | 30 | 0.253 | 0.180 | 0.714 |
| 8 – 38 species | 57 | 0.340 | 0.313 | 0.539 |
| 39 – 767 species | 27 | 0.464 | 0.435 | 0.435 |

These data also support a strong connection between pendant edge length and the FP index of evolutionary distinctiveness: fully 95 (>80%) have the highest-ranking FP species also being the species on the longest pendant edge. Importantly, this pattern is not driven by small-sized families; 26/30 clades with 3 to 8 species, 46/57 clades with 9 to 38 species, and 23/27 families with 39-767 species show this pattern.

We make no claims that taxa generally diversify under simple constant-rate birth-death processes - for instance, the full mammal phylogeny we use shows evidence consistent with multiple diversification rate shifts within Families (Upham et al., 2021). However, and interestingly, Morlon et al. (2010) found that fully 30% of 289 sampled phylogenies had shapes consistent with Yule expectations, and a further 35% had shapes consistent with a slowing diversification rate through time, an expectation consistent with both tree-inference biases (Revell et al., 2005) and speciation theory (see, e.g. Etienne and Rosindell (2012)). Taken together, this suggests that both real and expected trees are likely to contain particularly long pendant edges.

## Relevance for the amount of sequence data required to accurately infer a fully resolved tree

Both too-short and too-long edges can decrease the probability of correctly inferring a phylogenetic tree. In this section, we describe the impact of the interplay of long and short edges on the number of aligned DNA sequence sites required to accurately infer a phylogenetic tree from sequence data. The evolution of an aligned DNA sequence site is typically modelled by a continuous-time Markov process on a finite state space (typically the 4-element nucleotide set {*A, C, G, T*}) operating along the edges of the tree. Suppose that each site evolves independently along the edges of the tree with a fixed substitution rate *v* per site (where *μ* is constant across the edges and sites). We wish to consider the number of sites *K* required to infer *T* correctly with a given high probability.

Note that *K* depends on the tree, the edge lengths, the model of site substitution and the desired accuracy of tree reconstruction. *K* also depends on the tree reconstruction method applied, and so we consider here the usual form of maximum likelihood estimation without rate variation across sites (i.e. the substitution rate can vary across the tree but not across sites, and these along with the edge lengths are treated as nuisance parameters for the reconstruction). In particular, the inference of the tree does not include specifying the location of the root vertex.

It has been shown that for any binary phylogenetic tree with *n* leaves that has ‘carefully chosen’ edge lengths, *K* can grow as slowly as log(*n*) at least for simple site substitution models; a somewhat surprising and nontrivial result due to Daskalakis et al. (2011) (see also (Mossel et al., 2011; Mossel and Steel, 2004)). However, as noted by Felsenstein (2004) (pp. 173-174), edge lengths are likely to be variable and depend on the number of leaves of a tree (*n*) so a logarithmic dependence of *K* on *n* should lead to a faster growth function with *n*. Here, we describe a lower bound on *K* that is a positive power of *n* (rather than logarithmic in *n*), when the tree is generated by the Yule model (thus, there are now two random processes at play - the generation of the tree and its edge lengths, and the evolutions of sites on that tree).

### Proposition 3

Consider a Yule tree *T* grown for time *t* and a Markovian site-substitution model with site-substitution rate *v* operating on this tree. The sequence length *K* required to accurately reconstruct *T* from its associated sequence of sites at the leaves is bounded below by a term of order 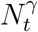 as λ*t* → ∞, where *γ* > 0 and *N_t_* is the number of leaves of the tree. Moreover, a lower bound of this type can be made independent of the value of *v* (for a possibly smaller but strictly positive value of *γ*).

The proof of Proposition 3, given in the Appendix, combines our earlier result on the longest pendant edge length in a Yule tree *T* with the following additional result concerning the length of the *shortest* interior edge in *T*.

### Proposition 4

Let 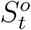 denote the length of the shortest interior edge in a Yule tree grown for time *t*.

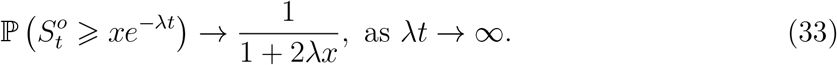

The proof of Proposition 4 is provided in the online Supplementary Material.

## Concluding comments

The simplest constant-rate birth-death process continues to offer up surprising tree shapes. While we know that any particular tree shape can be created under an infinite number of time-variant histories (Louca and Pennell, 2020), the constant-rate model is the default for species-level phylogenetic inference (e.g. in the popular RevBayes and BEAST packages); simple tree statistics based on the birth-death process (e.g. the methods of moments estimator of diversification rate; Magallon and Sanderson (2001)) have predictive power (see, e.g. Greenberg et al. (2021)); and limited surveys of the shapes of inferred trees and alternative models of diversification suggest the process we model here might underestimate pendant edge lengths in particular (viz. Morlon et al. (2010)). Remarkably old lineages (e.g. living fossils such as the tuatara, bichir, *Welwitschia* and *Amborella*) are well-known. The stochastic nature of speciation and extinction as we investigated here suggests some of these may (just) be expected statistical outliers (viz. Liow (2007)). And to the extent that the models capture the variation in edge lengths expected in large clades, we can expect our phylogenetic inferences to only slowly converge on the underlying true tree.

We also note that in the simulations (Fig. 3) as well as the mammal data (Table 1 and Fig. 4), trees with larger numbers of leaves tend to have larger values of 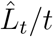 than trees with fewer leaves. In the theoretical results in our paper we are not conditioning on *n* (the number of leaves) apart from insisting that *n* ⩾ 1. Now, it might be expected that, for fixed times *t*, trees that have more leaves than the expected value (e.g. *e*^λ*t*^ for the Yule process) should have pendant edges that are slightly shorter on average, since the height of the tree is fixed, but the number of edges is increased (*cf*. Theorem 2 and Table 2 of Mooers et al. (2012)). However, our focus in this paper has been on the ‘longest’ pendant edge length (rather than the average), and increasing *n* may allow for more opportunity of an ‘outlier’ (extra-long) pendant edge to be present. For the mammal families, of course, *t* is not fully fixed and diversification rates vary - larger families may either have stochastically larger pushes of the past, or be governed by lower *μ* or higher λ*t* values.

**Fig. 4.**
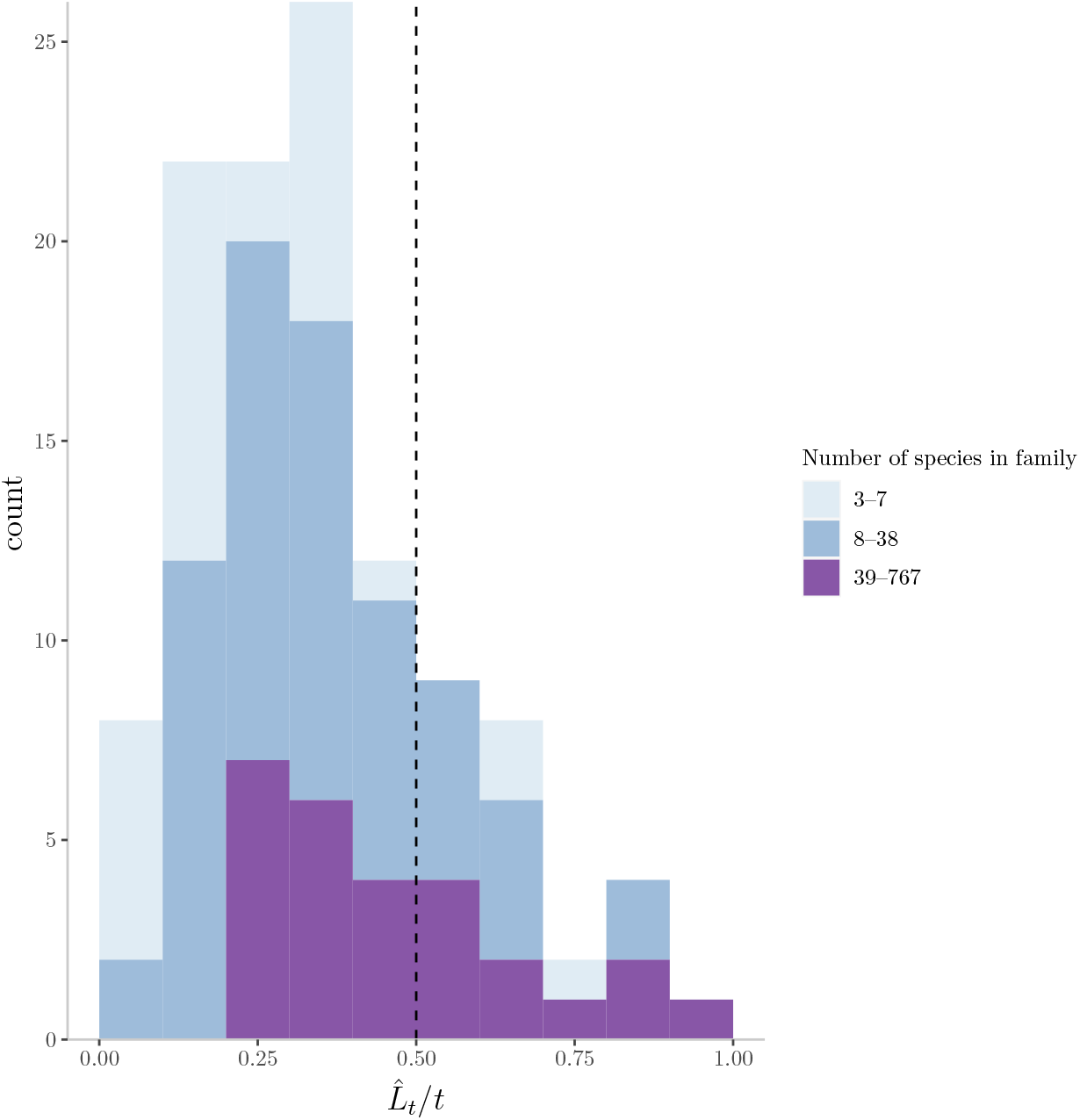
A histogram of the values 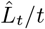 across 114 mammal families, broken up into the three groups (3-7 species, 8-38 species and 39+ species) distinguished by separate colours (stacked on each other) and with further statistical details provided in Table 1.

**Fig. 5.**
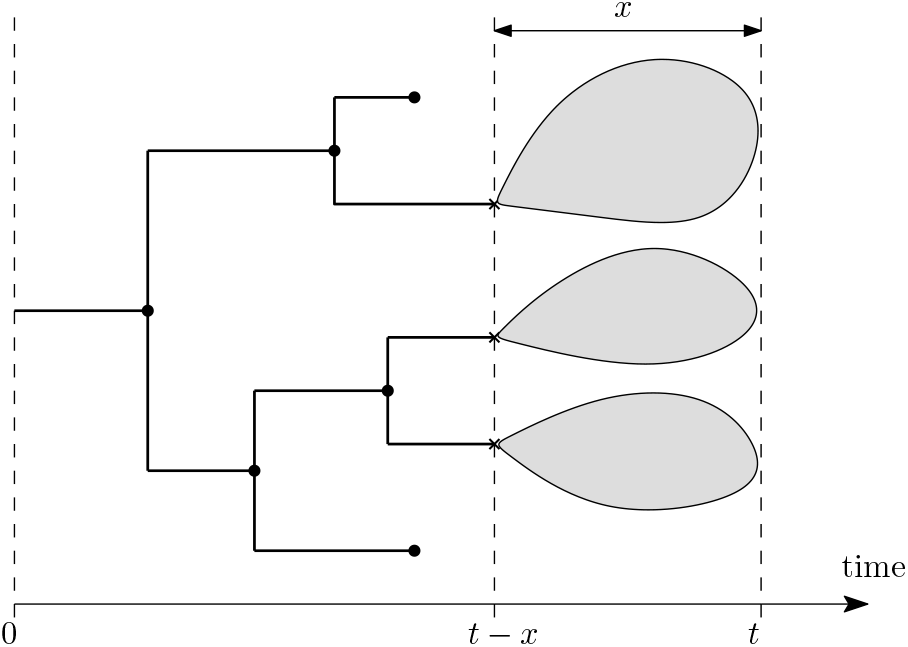
Subtrees of the birth–death tree 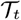 initiated at time *t* – *x* behave like *N_t–x_* independent birth–death trees each distributed like 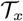. A pendant edge at time *t* of length greater than *x* must have been present at time *t – x* and not have any other birth or death event occur over remaining time of length *x*, that is, a subtree consisting only of a single edge.

Finally, it would be of interest to investigate the length of the longest (and shortest) edges in phylogenetic diversification models in which the birth and death rates (λ, *μ*) are not treated as constants, but allowed to depend on time *t*, or perhaps on other (stochastic) aspects of the branching process; for example the number of other lineages present in the tree at the given time, or the ‘age’ of the lineage (the time back to when it first split off from another lineage).

## Supplementary Material

The comparative data (mammal family trees) used in this study were retrieved from previously published phylogenies (Upham et al., 2018). A set of 500 global mammal phylogenies was downloaded from VertLife.org: http://vertlife.org/phylosubsets/. Additional mathematical results are available in the online Supplementary Material file.

## Acknowledgements

AOM and EK thank the Natural Sciences and Engineering Research Council (NSERC) Discovery Grant (AOM), NSERC PGSM scholarship (EK), and the NSERC CREATE program (‘RenewZoo’ training grant; AOM and EK). SB is supported by NSFC grant (No.11731012) and MS thanks the NZ Marsden Fund (MFP-UOC2005) for funding support. We thank Dr Paul Lewis and a second (anonymous) reviewer for several helpful comments and suggestions on an earlier version of this manuscript.

## Appendix: Mathematical details and proofs

### Part One: Yule tree

#### Proof of Proposition 2

We have:

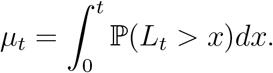

Thus, by Proposition 1 (Eqn. (1)):

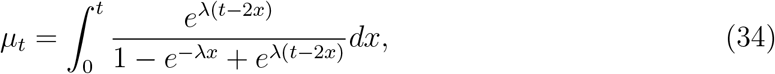

Making the substitution ζ = λ(*x* – *t*/2), this last integral can be written as:

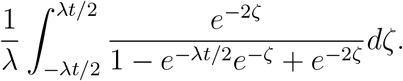

By making the further substitution *ξ* = *e*^−ζ^, we obtain:

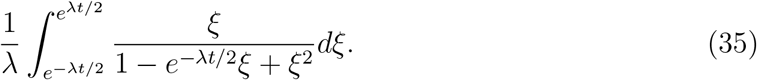

We now apply a standard integral result:

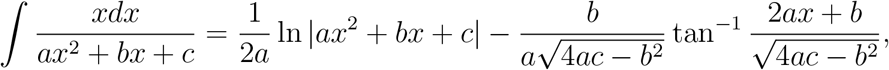

provided that 4*ac* – *b*^2^ > 0. Setting *a* = *c* = 1, *b* = – *e*^−λ*t*/2^ and *x* = *ξ*, Expression (35) becomes:

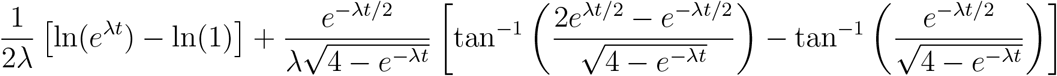

Finally, since 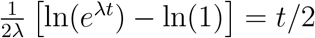, we obtain the claimed expression. The claimed limit as λ*t* → ∞ now follows.

### Part two: Complete birth–death tree

#### Proof of Lemma 2

Let *N_t_* be the number of lineages alive at time *t* in the birth-death process. By Lemma 1, *N_t_* has a *modified geometric distribution* on {0,1,2, 3,… }, where:

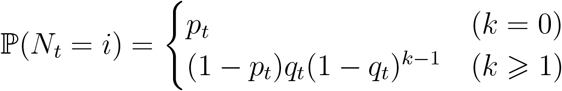

where *p_t_* and *q_t_* are as given in Lemma 1. We write *N_t_* ~ ModGeom(*p_t_, q_t_*). In particular, the average number alive at time *t* is 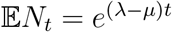, and the survival probability is

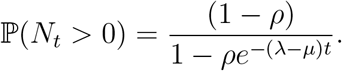

Note, the supercritical case corresponds to *ρ* ∈ [0,1) and in that case 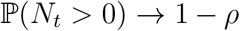 as *t* → ∞.

Observe that every pendant edge at time *t* that has length greater than *x* must have both been present at time *t* – *x* and also not have split into two lineages during the remaining time period (*t* – *x*, *t*]. Since each of the *N_t–x_* lineages alive at time *t* – *x* evolve forward in time independently, and (using the memoryless property of exponential lifetimes) each has an independent probability of success of *e*^−(λ+*μ*)*x*^ to give rise to a pendant edge of length at least *x* by not having any split or extinction event over the remaining time. Thus, conditional on *N_t–x_*, we have 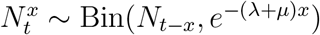.

A binomial random variable *Z* ~ Bin(*N, r*), where each of the *N* independent trials has a probability r of success but the total number of trials *N* ~ ModGeom(*p, q*) is an independent random variable, also gives rise to a modified geometric distribution for the total number of successes, where:

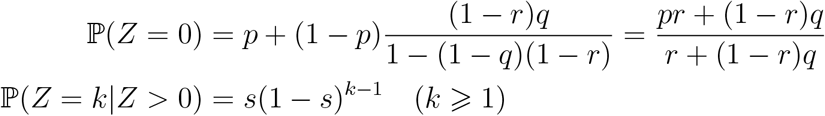

where *s* ≔ *q*/(1 – (1 – *q*)(1 – *r*)).

Simplifying these parameters above for the special case when 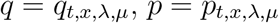, and *r* = *e*^−(λ+*μ*)*x*^ gives the claimed distribution for 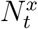.

#### Proof of Theorem 1

This result largely follows as a corollary to the explicit distribution for 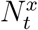 given in Lemma 2. Recalling the formulae for *p*_*t,x*,λ, *μ*_ and *q*_*t,x*,λ, *μ*_ given in Eqns. (8) and (9) respectively, setting 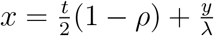 and then letting λ*t* → ∞, we find:

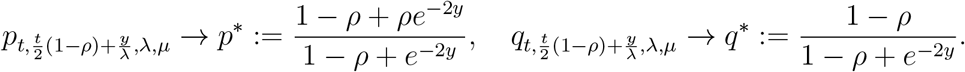

We also observe that (1 – *p*\*)/(1 – *ρ*) = 1 – *q*\* and recall that 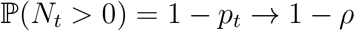 as λ*t* → ∞. Then, for *k* ⩾ 1:

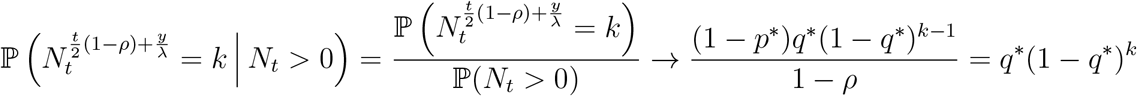

by using (10), conditional probability, and noting that *N_t_* > 0 is guaranteed whenever 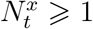. Similarly, also noting 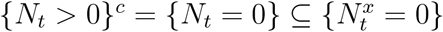, we find

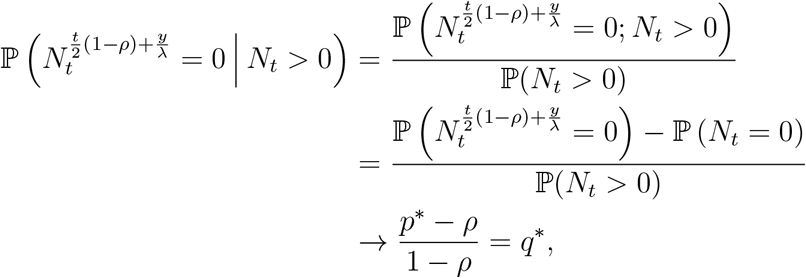

yielding Part (i). For Part (ii), we need simply note that:

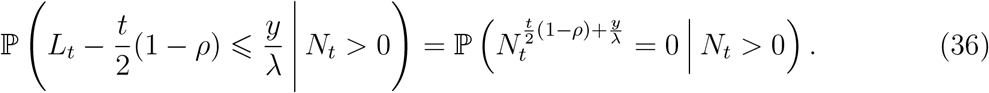

For Part (iii), we similarly observe that for *k* ⩾ 1,

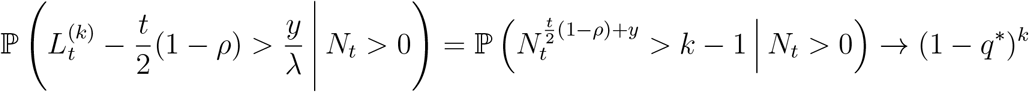

It remains to show the convergence in means. The arguments for Parts (ii)-(iii) are identical, so we discuss only (ii). As a simple consequence of (36), *L_t_/t* converges in distribution to the constant (1 – *ρ*)/2 whenever λ*t* → ∞, that is,

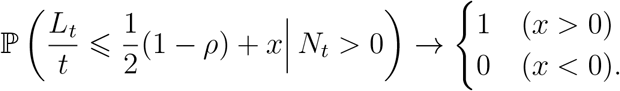

Now, convergence in distribution to a *constant* also implies convergence in probability. In other words, for all *ϵ* > 0:

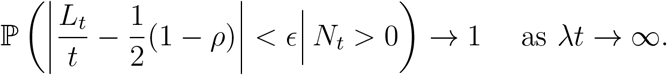

Finally, since *L_t_/t* ∈ [0,1], we can use the bounded convergence theorem to deduce that 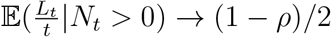 as λ*t* → ∞.

In addition to the convergence in means in the theorem above, we note that some finer almost sure convergence results also hold for 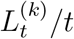. In particular, 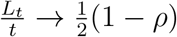 as *t* → ∞ almost surely (i.e. 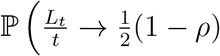 as 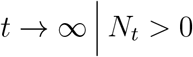 for all *t* ⩾ 0) = 1).

However, these are omitted as they would require a substantial additional analysis beyond the scope of the present article.

### Part 3: Reduced tree

#### Definition of 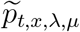 and 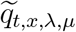

For *x* ∈ [0, *t*], let:

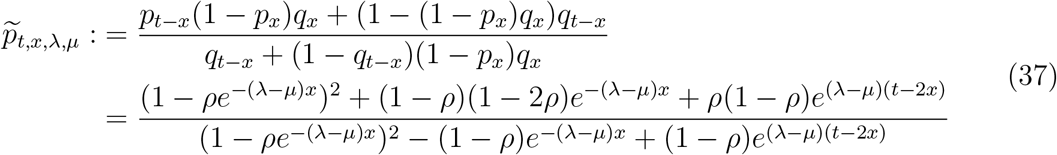

and

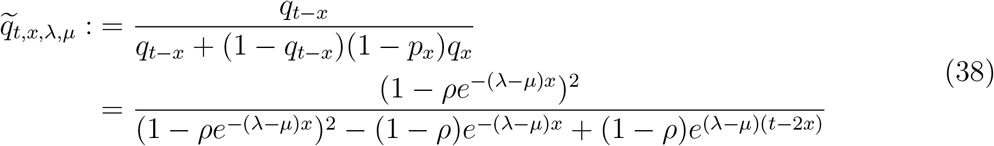

#### Proof of Lemma 3

We proceed by modifying the proof of Lemma 2. Firstly, recall that *N_t_* ~ ModGeom(*p_t_, q_t_*). Observe also that every pendant edge in the reduced tree at time *t* that has length greater than *x* must have come from some individual present at time *t* – *x* that has *exactly* one descendant alive after additional time *x* elapses. Now, each of the *N_t–x_* lineages alive at time *t* – *x* evolve forward in time independently, and each has a probability of success of 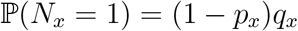 of giving rise to a pendant edge in the reduced tree at time *t* having length at least *x*. Thus, conditional on *N_t–x_*, we have 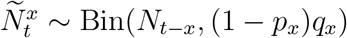.

In general, if *N* ~ ModGeom(*p, q*) and, conditional on *N*, we have *Z* ~ Bin(*N, r*), then we know that 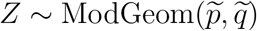, where:

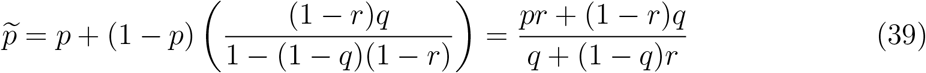

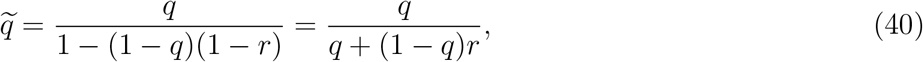

Simplifying these parameters above for the special case when *p* = *p_t–x_*, *q* = *q_t–x_* and *r* = (1 – *p_x_*)*q_x_* gives the claimed distribution for 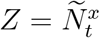.

#### Proof of Corollary 2

This follows directly from the distribution of 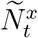 given in Lemma 3, since 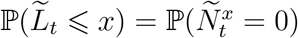 and 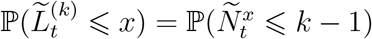.

#### Proof of Theorem 2

The proof of this result is essentially the same as the proof of Theorem 1 for the complete tree. It follows directly from the explicit distribution for 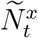 given in Lemma3 (instead of Lemma2 in the complete tree analogue), combined with the limits given in (24), (25), (23), together with the simple observation that 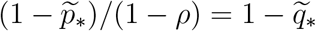.

### Proof of Eqn. (32)

Let 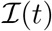 (resp. 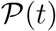) denote the sum of the interior edges (resp. pendant edges) in a Yule tree grown for time *t*. Summing over the leaves of the tree that are present at time *t* we have 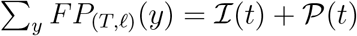 (since the sum of *FP* over all leaves is the total sum of edge lengths in the tree) and 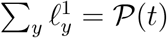. By Theorem 4 of Steel and Mooers (2010), 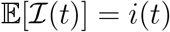 and 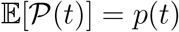, where 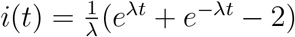 and 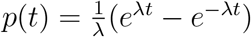. Observe that 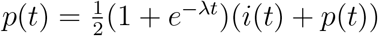, and so

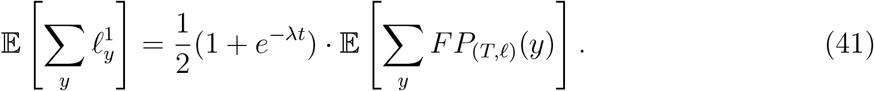

Now consider a leaf *x* selected uniformly at random from the *N* leaves present at time *t* (note that *N* is a random variable). We have 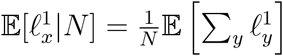 and 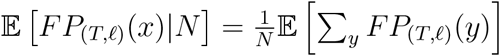, and so, from Eqn. (41),

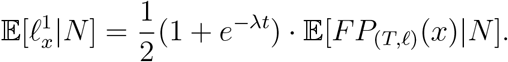

This equality holds for all values of *N* and so taking a further expectation (over *N*) yields Eqn. (32).

### Proof of Theorem 3

The proof of Theorem 3 relies on the following two claims

i. With probability tending to 1 as λ*t* grows (with *ρ* = *μ*/λ fixed), every leaf *x* of the reduced birth-death tree *T* that has incident pendant edge length 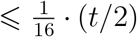 satisfies the inequality: 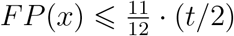.
ii. Let *δ* ∈ (0,1) be fixed, and consider a leaf *x* of the reduced birth-death tree *T* for which the incident pendant edge has length 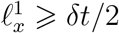. Then,

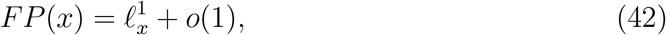

where *o*(1) is a positive term that converges in probability to 0 as λ*t* grows (with *ρ* = *μ*/λ fixed).

The proofs of Claims (i) and (ii) are provided shortly. First, we show how Theorem 3 follows from them. We may assume, without loss of generality, that 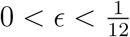. Let *e*′ be a longest pendant edge in *T* and let *x*′ be its end leaf. By Theorem 2, *e*′ has length at least (1 – *ϵ*)(*t*/2) with probability 1 – *o*(1) (as *t* grows), and since 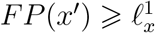, it follows that 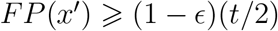 with probability 1 – *o*(1).

Next, consider any leaf *x* of *T*. If 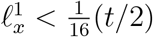 then, by Claim (i), 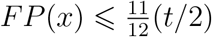 with probability 1 – *o*(1), and since 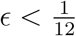 it follows that *FP*(*x*) < *FP*(*x*′) so *x* cannot be a leaf that maximises *FP*. Thus any leaf that maximises *FP* satisfies ℓ_*x*_ ⩾ *δ*(*t*/2) for 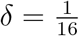. It then follows from Claim (ii) that (for any such leaf *x* satisfying this last inequality) we have: 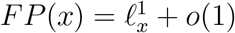, and so any leaf with maximal FP value will have 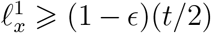 as λ*t* grows (such a leaf *x*′ exists as shown at the start of this proof). This completes the proof of Theorem 3 modulo verifying the two claims.

**Proof of Claim (i):** We first state a lemma, the proof of which is provided in the online Supplementary Material.

#### Lemma 4

For any *ϵ* > 0, the probability the reconstructed birth-death tree has an interior edges of length greater than 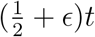 decays exponentially fast to 0 as λ*t* → ∞ (with *ρ* = *μ*/λ fixed).

Next, observe that for any leaf *x*, 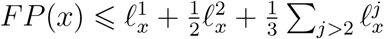, and since 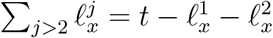 it follows that 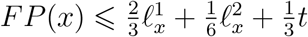. In particular, if 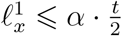 and 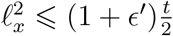 then:

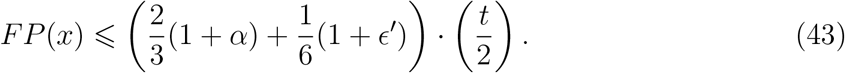

Note that 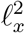 is an interior edge, and so the condition that 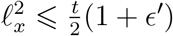 holds with probability 1 – *o*(1) by Lemma 4. Taking *α* = 1/16 and *ϵ*′ = 1/4 and we see that the right-hand side Inequality (43) is 11/12, which establishes Claim (i).

**Proof of Claim (ii):** Notice that if *n*_2_(*x*) ⩾ *M*, then:

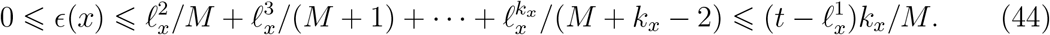

Suppose that *x* is a tip of a pendant edge *e* of a reduced birth-death tree *T* grown for time *t*, and that *e* has length ℓ_*x*_ ⩾ *δt*/2 for some constant *δ* > 0. Consider the subtree of *T* descending from the other endpoint of edge *e*_1_ to *x*. By the assumption that *T* has evolved according to a birth-death model, the tree topology is described by the Yule-Harding model, and thus the number of leaves of this subtree (i.e. *n*_2_(*x*) – 1) is geometrically distributed with parameter 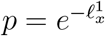. In particular, since 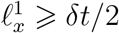, it follows that *n*_2_(*x*) is at least *e*^(λ–*μ*)*δt*/4^ probability converging to 1 as *t* grows. Furthermore, since the topology of a reduced birth-death tree with *n* leaves is described by the Yule-Harding distribution (i.e. the *β*-splitting model with *β* = 0) the number of edges on the path from the root to a most distant leaf is concentrated around a term of order log(*n*) (Proposition 4 of Aldous (1996)) and thus *k_x_* is of order log(*n*) which, by Jensen’s inequality, grows at most linearly in expectation with λ*t*. Consequently, from Inequality (44), *ϵ*(*x*) is (with high probability as λ*t* grows) bounded above by a term of order λ*t*^2^*e*^−λ*t*/4^, and so for an edge of length ⩾ *δt*/2,

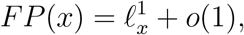

where *o*(1) refers to a positive term that converges in probability to 0 as *t* grows. This establishes Claim (ii).

We end this section by noting that for a randomly selected extant species *x* in a reconstructed birth–death tree, the equation 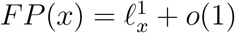 (Eqn. 42) fails to hold. This is because the expected value of 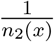 for the Yule-Harding distribution (e.g. a birth-death tree) with *n* leaves is given (via Eqn. (3.8) in Steel (2016)) by:

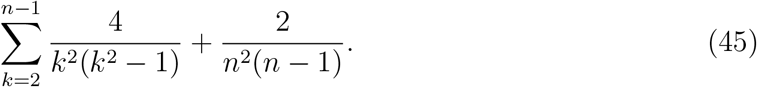

By using partial fractions, and the identity 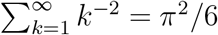, the expression in (45) simplifies to *β* + *o*(1), where *β* = 7 – 2*π*^2^/3 ≈ 0.42. Thus, up to a term of vanishing order *o*(1), the expected value of *FP*(*x*) for such a tree (where expectation refers to the tree shape produced by the birth-death process) is: 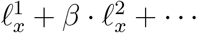. Thus, for a randomly selected species *x*, the size of 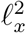 has a non-vanishing influence on the FP score of *x*.

### Proof of Proposition 3

Given a fully-resolved (i.e. binary) tree *T* with edge lengths, let ℓ_+_ denote the length of the longest pendant edge and let ℓ_−_ denote the length of the shortest interior edge (we use ℓ_+_ rather than *L_t_* to distinguish between actual edge lengths on a given tree versus (random) edge lengths for a tree generated by a model).

The proof of Proposition 3 combines two results:

i. *K* is bounded below by a term that grows at the rate *e^cνℓ_+_^* as ℓ_+_ → ∞, and
ii. *K* is bounded below by a term of order 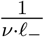 as ℓ_−_ → 0.

(for details, see Section 8.2.1 of Steel (2016); note that in the constant c is strictly positive and depends (only) on the particular model of site substitution). The intuition behind Result (i) is that for a very long edge *e*, incident with leaf *x*, sites evolved along *e* are likely to have undergone multiple substitutions and this erases the signal in the data concerning where leaf *x* attaches to the rest of the tree. The intuition behind Result (ii) is that if none of the sites in the data have evolved with a substitution on internal edge *e* then *e* cannot be detected from the data.

By Proposition 1(iii), ℓ_+_ ~ *t*/2 as λ*t* grows. Consequently, Result (i) from the previous paragraph implies that *K* is bounded below by *e*^*cνt*/2^ as λ*t* → ∞. Since *N_t_* has a geometric distribution with mean *e*^λ*t*^, as λ*t* → ∞:

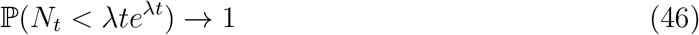

So, with probability 1 as λ*t* → ∞,

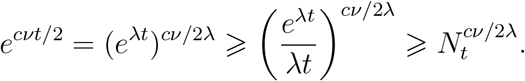

To explore the impact of Result (ii) we take *x* = *t* in Proposition 4 to obtain 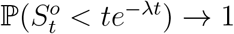 as λ*t* → ∞, and so Result (ii) implies that *K* is bounded below by a term of order *e*^λ*t*^/(*vt*) as λ*t* grows. Moreover, by Eqn. (46), and any *ϵ* > 0, with probability tending to 1 as λ*t* → ∞:

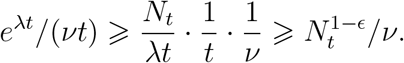

Comparing these two lower bounds on *K* (namely, 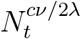 and 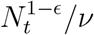) reveals that if *v* ⩾ 1 then we can take *γ* = *c*/2λ, while if *v* < 1 we can take *γ* =1 – *ϵ* (for any *ϵ* ∈ (0,1)), justifying the second claim of a lower bound on *K* that is a positive power of *N_t_* and independent of *v*.

## SUPPLEMENTARY MATERIAL

**Abstract**

We provide some supplementary material for the article [1] that concerns (a) shortest interior edges in Yule trees (proving Proposition 4 in [1]), and (b) long interior edges in time inhomogeneous Yule trees (which includes reconstructed Birth-Death trees, proving Lemma 4 in [1]). Throughout, we will use models and notation given in [1].

### 1 Short interior edges

#### 1.1 Shortest interior edge of a Yule process

Let 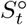 be the *shortest interior edge* up to time *t* in the Yule process with birth rate λ, that is, the shortest of all edges excluding any pendant edges that are alive at time *t*. Then, 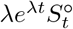 converges in distribution to *W* where

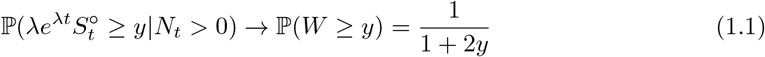

as λ*t* → ∞.

*Proof of* (1.1). A rigorous proof for the shortest interior edge in the Yule tree can be obtained by calculating the first and second moments of the number of short interior edges at time *t* of length less than 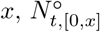. This will requires some quite involved calculations.

Recall that, in the Yule process, we have binary branching at rate λ (and no deaths). We fix an edge length *x* > 0. For times *t* > 0, we define the number of edges of length at most *x* given by

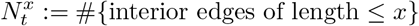

We will make good use of some asymptotics for the first and second moments (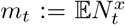 and 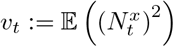)as in the following result:

##### Proposition 1.1.

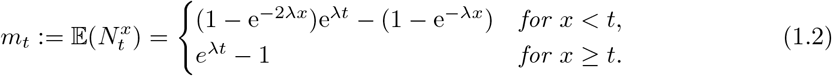

*In particular, whenever λt* → ∞ *and λx* → 0, *we have m_t_* ~ 2λ*xe*^λ*t*^. *Further, we find that the first and second moments are asymptotically equivalent, that is*,

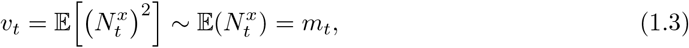

*whenever x depends on t such that x* = *o*(e^−λ*t*^) *as* λ*t* → ∞.

We prove this result shortly, but first we show how it can be used to establish the main result of this section, namely Eqn. (1.1).

Let us now fix any *y* > 0 and *δ* ∈ (0,1) and throughout we fix *x* ≔ *ye*^−λ*t*^ (in this way, *x* = *o*(e^−λ(1–*δ*)*t*^)). We let 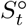 be the length of the shortest interior edge at time *t* so that

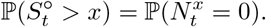

We consider the events

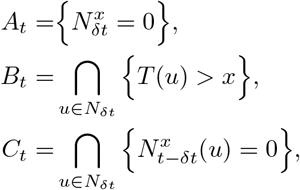

where for every *u* ∈ *N_δt_*, *T*(*u*) is the first branching time in the subtree initiated by particle *u* at time *δt*, and 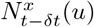 is the size of the population after time *t* – *δt* in the subtree initiated by particle *u* at time *δt*. We observe that

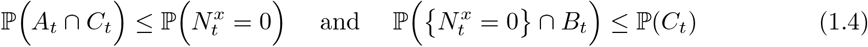

(Note, any edges that span time *δt* will be considered as two shorter edges in terms of events *A_t_* and *C_t_*, and event *B_t_* implies none of the edges alive at time *δt* can be shorter than *x*.) The idea is to show that, as λ*t* → ∞, 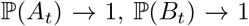, and 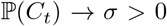 where we can readily calculate this limit, and hence, by the bounds in (1.4), this necessarily implies 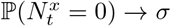.

To that end, since

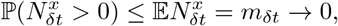

it follows that

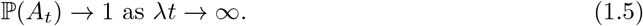

Also, since

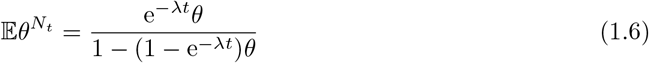

and as subtrees initiated by particles *u* ∈ *N_δt_* are independent copies of the original tree, it follows that

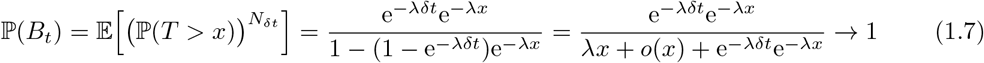

as λ*t* → ∞ (recall, *x* = *ye*^−λ*t*^ ≪ e^−λ*δt*^).

Finally, to estimate 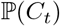, recalling the moment asymptotics of Proposition 1.1, we note that

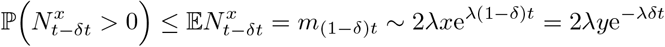

and, using the Paley-Zygmund inequality,

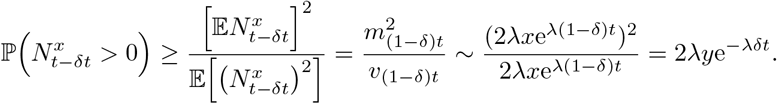

Thus, these asymptotic upper and lower bounds agree, so we have

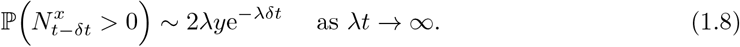

Then, using (1.6) again, we find that

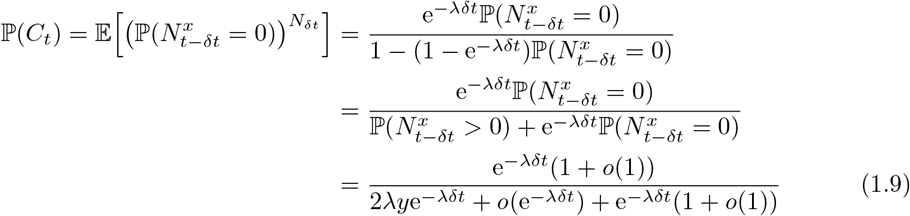

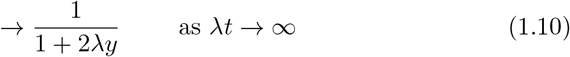

In conclusion, combining (1.5), (1.7) and (1.9) with inequalities (1.4), we find

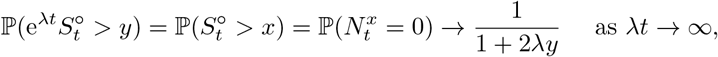

as required.

It only remains to confirm the moment results for the number of interior edges less than length *x* at time *t*.

*Proof of Proposition 1.1. (a) First moment*. If *t* ∈ (0, *x*], then every interior edge is ≤ *x*. That is, 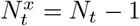, where *N_t_* is the population size at time *t*. Thus

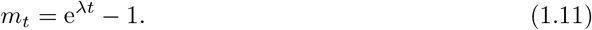

If *t* > *x*, then by considering when the first birth *T* ~ Exp(λ) occurs, we have

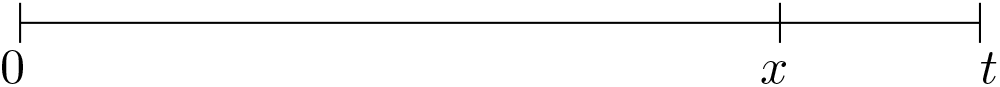

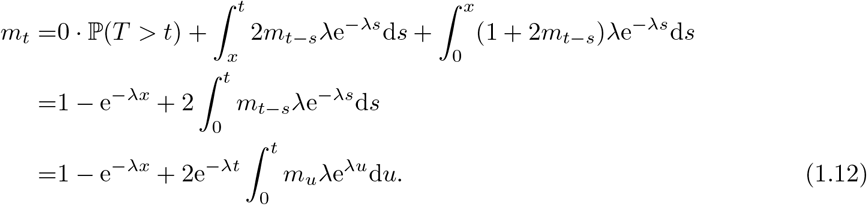

From (1.12) and (1.11) it follows that

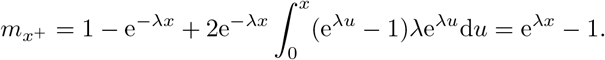

Furthermore, from the integral equation (1.12) we see that *m* is continuous and differentiable on (*x*, ∞). Differentiating (1.11) yields

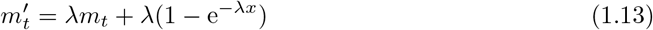

Multiplying (1.13) by the integrating factor e^−λ*t*^ gives

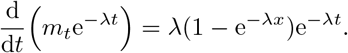

Integrating the above equation between *x* and *t* gives

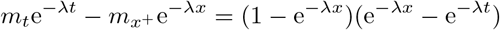

and hence

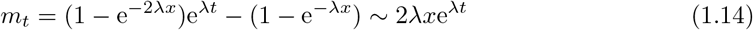

whenever λ*x* → 0 and λ*t* → ∞ since *e^y^* = 1 + *y* + *o*(*y*) for *y* → 0.

(*b) Second moment*. If *t* ∈ (0, *x*] then, as before, every interior edge is at most length *x*. That is, 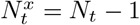 and hence

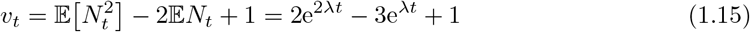

If *t* > *x*, then once again by considering when the first birth *T* ~ Exp(λ) occurs, we find

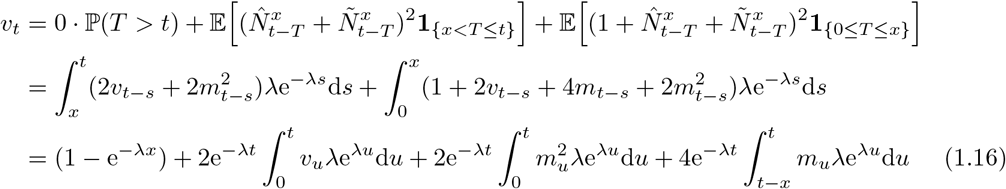

In particular, *v_t_* is continuous on (*x*, ∞) and hence also differentiable on (*x*, ∞). We may also find from the integral equation (1.16) and also (1.11) and (1.15) that

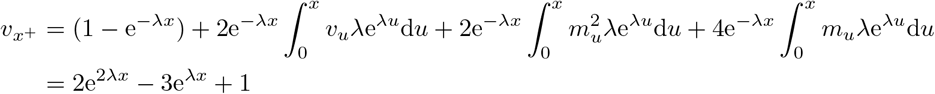

In particular, *v_x_*+ ~ λ*x* as *x* → 0. In order to help find its solution, the integral equation (1.16) can be converted to the following differential equation

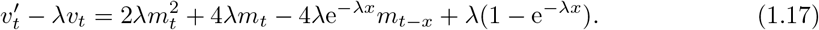

Equivalently,

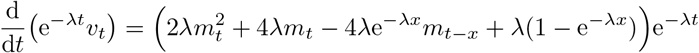

which needs to be solved separately in intervals (*x*, 2*x*) and (2*x*, ∞) due to the fact that *m_t–x_* has a different form in each of these two intervals. We find that

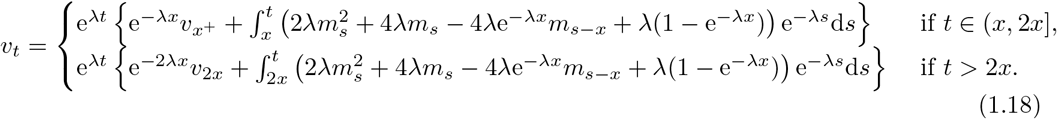

Rather than evaluating this exactly, which is unnecessary for our purposes, we find from the first line of (1.18) that

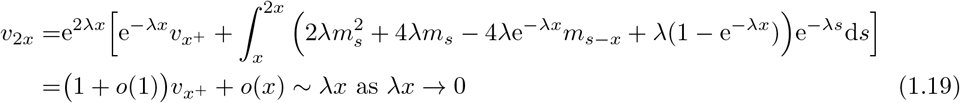

(In the integral, one should use (1.14) for *m_s_* and (1.11) for *m_s–x_.*)

The second line of (1.18) says that

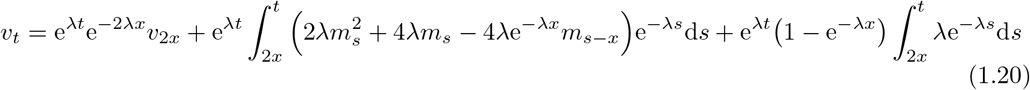

If we now choose *x* such that *x* = *o*(e^−λ*t*^) as λ*t* → ∞ and use (1.19) for *v*_2*x*_, (1.14) for terms *m_s_* and *m_s–x_* inside the integral then (1.20) becomes

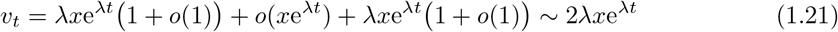

as required.

### 2 Long interior edges in time inhomogeneous Yule processes

Consider a *time inhomogeneous Yule process* where each individual alive at time *s* independently splits into 2 individuals at (time dependent) instantaneous rate λ_s_. That is, for example, for small times *h* > 0,

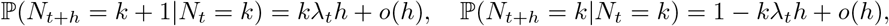

and the probability there has been no birth event by time *t* is given by

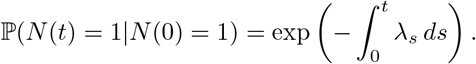

We will assume throughout that λ_*s*_ is a continuous function in time parameter *s*.

#### Example 2.1

*The following are special cases of time inhomogeneous Yule processes*:

i. *The classical Yule process with constant branching rate has* λ*_s_* ≡ λ.
ii. *The reconstructed (or reduced) tree derived from the birth-death process with birth rate* λ *and death rate μ has instantaneous branching rate*

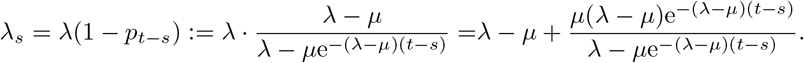 *We can interpret this rate intuitively as follows. Recall, any individual that remains in the reduced tree must have at least one descendant still alive at time t (ie. any family lines that go extinct before time t are removed). Now, any individual alive at time *s* that is part of the reduced tree at time t will give birth to another individual at rate* λ *but this individual will only be in the reduced tree if its descendants survive to time t from its birth at time *s*, where this has probability* 1 – *p_t–s_ (as defined above). Thus, the effective rate of births in the reduced tree becomes* λ(1 – *p_t–s_*).

We wish to find the expected number of interior edges in an inhomogeneous Yule process at time *t* that are of length at least *x* when starting from a single ancestor at time 0.

- Fix a time *t* > 0 and an edge length 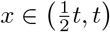.
- For any time *τ* ∈ [0, *t* – *x*] we let 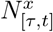 be the number of interior edges of length at least *x* at time *t* in Yule process with instantaneous branching rate λ*_s_* initiated at time *τ*. (Here, interior edges at time *t* means all edges up to time *t* except for pendant edges alive at *t*.)
- Define 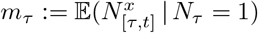
- Note, whenever *t* > 0, *x* ∈ (*t*/2/*t*), and *τ* ∈ [0, *t* – *x*], we have *τ* ≤ *t* – *x* ≤ *τ* + *x* ≤ *t*.

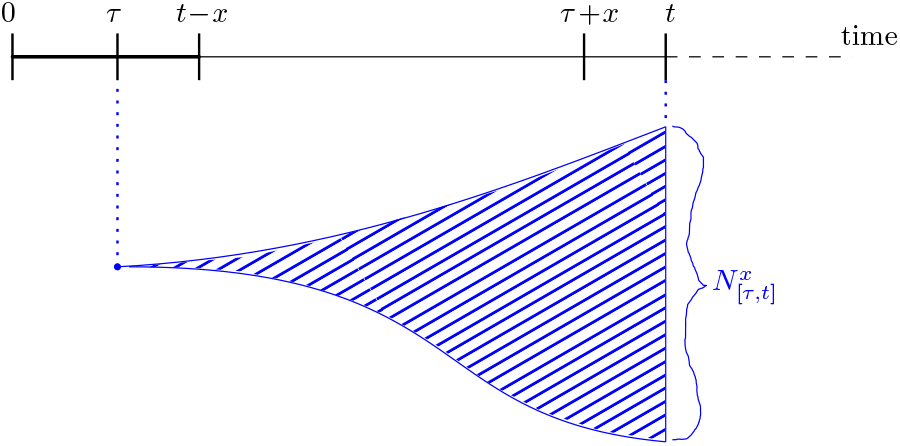

#### Lemma 2.2.

*For t* > 0 *and x* ∈ (*t*/2, *t*),

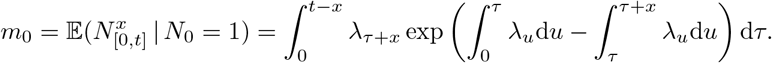

*Proof of Lemma 2.2*. Let *T* be the first branching time in the process initiated from a single particle at time *τ*. Then, for *s* > *τ* ≥ 0,

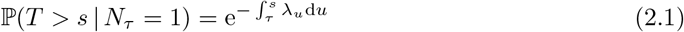

and we find that *m_τ_* satisfies the following integral equation

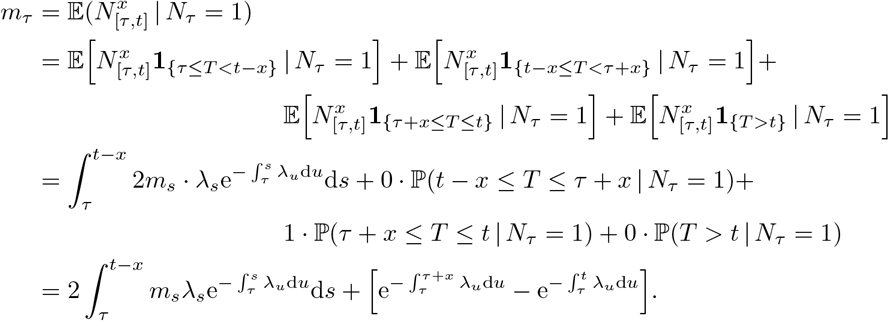

In the above decomposition, the first term covers the first birth event occurring at any times *s* ≤ *t* – *x*, so the first interior edge is not longer that *x* (ie. too short to be counted) and we add up edges from the 2 independent offspring starting at time *s*. The second term covers branching events at times *s* ∈ [*t* – *x*, *τ* + *x*], so this first interior edge is shorter than *x* (ie. no contribution as occurs too early before time *τ* + *x*) and neither offspring can result in any interior edges longer than *x* (since they start too late after time *t* – *x*). In the third term, the first birth event occurs at time *s* ∈ [*τ* + *x*, *t*] so this first interior edge is longer than *x* but the offspring cannot have any interior edges longer than *x*, hence exactly 1 interior edge results. Finally, the last term covers when there is no birth event before time *t*, hence no interior edges at all.

Differentiating this integral equation with respect to *τ* and also taking *τ* = *t* – *x* for the initial condition (where the first birth would have to occur *exactly* at time *t* to give 1 interior edge of length at least *x*, and this has zero probability), we derive the following first-order linear ordinary differential equation.

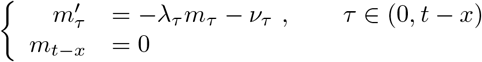

where

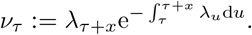

To solve this differential equation we multiply it by the integrating factor 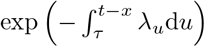, so

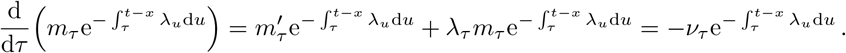

Then integrating over *τ* from 0 to *t* – *x* reveals that

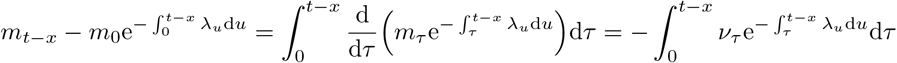

Finally, using boundary condition *m_t–x_* = 0, we obtain

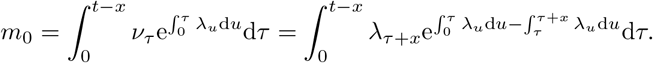

#### Example 2.3

(Classical Yule process). *In the special case of constant branching where* λ*_s_* ≡ λ,

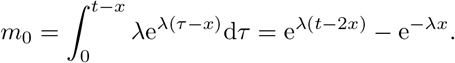

#### Corollary 2.4

(Atypically long interior edges in reconstructed birth-death tree). *For some constant C and any sufficiently small ϵ* > 0,

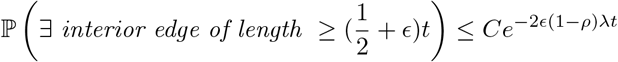

*That is, the probability of finding an interior edge longer than* (1/2 + *ϵ*)*t in the reconstructed birth-death tree decays exponentially fast to zero as* λ*t* → ∞ (*where ρ* ≔ *μ*/λ *is fixed*)

*Proof of Corollary 2.4*. It is known that the reduced (reconstructed) tree of the birth-death process with birth rate λ and death rate *μ* is a time inhomogeneous Yule tree with instantaneous branching rate

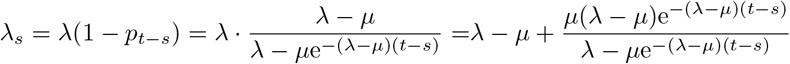

We note that for all *s*,

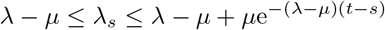

and trivially λ*_s_* ≤ λ. Combining these simple bounds with Lemma 2.2, we find

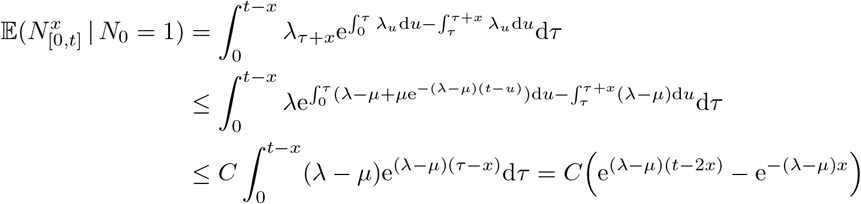

for some constant *C*.

Thus, for small *ϵ* > 0, we have 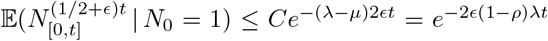. Further, by applying Markov’s inequality,

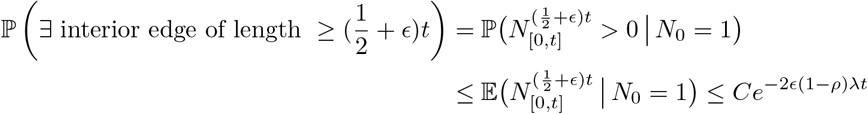

which is decaying exponentially fast to zero as λ*t* → ∞ (where *ρ* = *μ*/λ is fixed), as required.

## References

Aldous, D. 1996. Probability distributions on cladograms. Pages 1–18 in Random Discrete Processes (Eds. D Aldous and R. Pemantle) vol. 76 of IMA volumes in Mathematics and its Applications. Springer.

Aldous, D. and L. Popovic. 2005. A critical branching process model for biodiversity. Adv. Appl. Probab. 37:1094–1115.

Aldous, D. J. 2001. Stochastic models and descriptive statistics for phylogenetic trees, from Yule to today. Statist. Sci. 16:23–34.

Burgin, C. J., J. P. Colella, P. L. Kahn, and N. S. Upham. 2018. How many species of mammals are there? J. Mammal. 99:1–14.

Daskalakis, C., E. Mossel, and S. Roch. 2011. Evolutionary trees and the Ising model on the Bethe Lattice: A proof of Steel’s conjecture. Prob. Theor. Rel. Fields 149:149–189.

Etienne, R. S. and J. Rosindell. 2012. Prolonging the past counteracts the pull of the present: Protracted speciation can explain observed slowdowns in diversification. Systematic Biology 61:204–213.

Felsenstein, J. 2004. Inferring Phylogenies. Sinauer Associates, Sunderland, MA.

Fuchs, M. and E. Y. Jin. 2015. Equality of Shapley value and fair proportion index in phylogenetic trees. J. Math. Biol. 71:1133–1147.

Gascuel, O. and M. Steel. 2010. Inferring ancestral sequences in taxon-rich phylogenies. Math. Biosci. 227:125–135.

Gavrilets, S. and A. Vose. 2005. Dynamics of adaptive radiation. Proc. Natl. Acad. Sci. U.S.A. 102:18040–18045.

Greenberg, D. A., R. A. Pyron, L. G. W. Johnson, N. S. Upham, W. Jetz, and A. O. Mooers. 2021. Evolutionary legacies in contemporary tetrapod imperilment. Ecol. Lett. 24:2464–2476.

Grimmett, G. and D. Stirzaker. 2001. Probability and Random Processes (3rd ed.). Oxford University Press.

Hey, J. 1992. Using phylogenetic trees to study speciation and extinction. Evolution 46:627–640.

Isaac, N. J. B., S. T. Turvey, B. Collen, C. Waterman, and J. E. M. Baillie. 2007. Mammals on the EDGE: Conservation priorities based on threat and phylogeny. PLoS ONE 2:e296.

Kembel, S. W., P. D. Cowan, M. R. Helmus, W. K. Cornwell, H. Morlon, D. D. Ackerly, S. P. Blomberg, and C. O. Webb. 2010. Picante: R tools for integrating phylogenies and ecology. Bioinformatics 26:1463–1464.

Kendall, D. G. 1948. On the generalized ‘birth-and-death’ process. Ann. Math. Statist. 19:1–15.

Lambert, A. and T. Stadler. 2013. Birth–death models and coalescent point processes: the shape and probability of reconstructed phylogenies. Theor. Popul. Biol. 90:113–128.

Liow, L. H. 2007. Lineages with long durations are old and morphologically average: an analysis using multiple datasets. Evolution 61:885–901.

Louca, S. and M. W. Pennell. 2020. Extant timetrees are consistent with a myriad of diversification histories. Nature 580:502–505.

Magallon, S. and M. J. Sanderson. 2001. Absolute diversification rates in angiosperm clades. Evolution 55:1762–1780.

Mooers, A. O., O. Gascuel, T. Stadler, H. Li, and M. Steel. 2012. Branch lengths on birth–death trees and the expected loss of phylogenetic diversity. Syst. Biol. 61:195–203.

Morlon, H., M. D. Potts, and J. B. Plotkin. 2010. Inferring the dynamics of diversification: a coalescent approach. PLOS Biol. 8:e1000493.

Mossel, E., S. Roch, and A. Sly. 2011. On the inference of large phylogenies with long branches: How long is too long? Bull. Math. Biol. 73:1627–1644.

Mossel, E. and M. Steel. 2004. A phase transition for a random cluster model on phylogenetic trees. Math. Biosci. 187:189–203.

Nee, S., E. C. Holmes, R. M. May, and P. H. Harvey. 1994. Extinction rates can be estimated from molecular phylogenies. Philos. Trans. R. Soc. Lond. B Biol. Sci. 344:77–82.

Paradis, E. and K. Schliep. 2019. ape 5.0: an environment for modern phylogenetics and evolutionary analyses and evolutionary analyses in R. Bioinformatics 35:526–528.

Pennell, M. W., J. M. Eastman, G. J. Slater, J. W. Brown, J. C. Uyeda, R. G. FitzJohn, M. E. Alfaro, and L. J. Harmon. 2014. geiger v2.0: an expanded suite of methods for fitting macroevolutionary models to phylogenetic trees. Bioinformatics 30:2216–2218.

Phillimore, A. B. and T. D. Price. 2008. Density-dependent cladogenesis in birds. PLOS Biol. 6:483–489.

Redding, D. W. 2003. Incorporating genetic distinctness and reserve occupancy into a conservation prioritisation approach. Masters Thesis, University Of East Anglia, Norwich, UK.

Redding, D. W., K. Hartmann, A. Mimoto, D. Bokal, M. Devos, and A. O. Mooers. 2008. Evolutionarily distinctive species often capture more phylogenetic diversity than expected. J. Theor. Biol. 251:606–615.

Revell, L. J. 2012. phytools: an R package for phylogenetic comparative biology (and other things). Methods Ecol. Evol. 3:217–223.

Revell, L. J., L. J. Harmon, and R. E. Glor. 2005. Underparameterized model of sequence evolution leads to bias in the estimation of diversification rates from molecular phylogenies. Systematic Biology 54:973–983.

Stadler, T. 2009. On incomplete sampling under birth-death models and connections to the sampling-based coalescent. J. Theor. Biol. 261:58–66.

Stadler, T. 2011. Simulating trees with a fixed number of extant species. Syst. Biol 60:676–684.

Stadler, T. and M. Steel. 2012. Distribution of branch lengths and phylogenetic diversity under homogeneous speciation models. J. Theor. Biol. 297:33–40.

Steel, M. 2016. Phylogeny: Discrete and Random Processes in Evolution. Society for Industrial and Applied Mathematics, Philadelphia PA.

Steel, M. and A. Mooers. 2010. The expected length of pendant and interior edges of a Yule tree. Appl. Math. Lett. 23:1315–1319.

Upham, N. S., J. A. Esselstyn, and W. Jetz. 2018. Inferring the mammal tree: Species-level sets of phylogenies for questions in ecology, evolution, and conservation. PLoS Biol. 17:e3000494.

Upham, N. S., J. A. Esselstyn, and W. Jetz. 2021. Molecules and fossils tell distinct yet complementary stories of mammal diversification. Curr. Biol. 31:4195–4206.e3.

Wicke, K., A. Mooers, and M. Steel. 2021. Formal links between feature diversity and phylogenetic diversity. Syst. Biol. 70:480–490.

Yule, G. U. 1925. A mathematical theory of evolution: Based on the conclusions of Dr. J.C. Willis F.R.S. Philos. Trans. R. Soc. Lond. B 213:21–87.

## References

[1] Sergey Bocharov, Simon Harris, Emma Kominek, Arne Ø. Mooers, and Mike Steel. Predicting long pendant edges in model phylogenies, with applications to biodiversity and tree inference. (2022)

